# *APOE* modulates microglial immunometabolism in response to age, amyloid pathology, and inflammatory challenge

**DOI:** 10.1101/2022.05.17.492361

**Authors:** Sangderk Lee, Nicholas A. Devanney, Lesley R. Golden, Cathryn T. Smith, James L. Schwarz, Adeline E. Walsh, Harrison A. Clarke, Danielle S. Goulding, Elizabeth J. Allenger, Gabriella Morillo-Segovia, Cassi M. Friday, Amy A. Gorman, Tara R. Hawkinson, Steven M. MacLean, Holden C. Williams, Ramon C. Sun, Josh M. Morganti, Lance A. Johnson

## Abstract

The E4 allele of Apolipoprotein E (*APOE*) is associated with both metabolic dysfunction and a heightened pro-inflammatory response – two findings that may be intrinsically linked through the concept of immunometabolism. Here, we combined bulk, single-cell, and spatial transcriptomics with cell-specific and spatially resolved metabolic analyses to systematically address the role of *APOE* across age, neuroinflammation, and AD pathology. RNAseq highlighted immunometabolic changes across the *APOE4* glial transcriptome, specifically in subsets of metabolically distinct microglia enriched in the E4 brain during aging or following an inflammatory challenge. E4 microglia display increased *Hif1α* expression, a disrupted TCA cycle, and are inherently pro-glycolytic, while spatial transcriptomics and MALDI mass spectrometry imaging highlight an E4-specific response to amyloid that is characterized by widespread alterations in lipid metabolism. Taken together, our findings emphasize a central role for *APOE* in regulating microglial immunometabolism.

## Main

Metabolic dysfunction and chronic neuroinflammation are two features common to several neurodegenerative diseases, including Alzheimer’s disease (AD). Top hits from genome-wide association studies indicate that the microglial immune response is central to AD risk^1–5^. Likewise, altered patterns of glucose and lipid metabolism are early biomarkers of incipient AD^6^, with proteomic and metabolomic studies strongly linking changes in glial glucose metabolism to cognitive impairment and AD pathology^7–10^. Tying metabolic dysfunction and neuroinflammation together is a well-established process whereby innate immune responses invoke metabolic reprogramming in microglia and vice versa^11,12^. However, it remains unclear how this phenomenon of immunometabolism may relate to AD etiology and genetic risk factors.

Intriguingly, the strongest genetic risk factor for late-onset AD, the ε4 allele of Apolipoprotein E (*APOE*), has been separately linked to both heightened neuroinflammation and alterations in glial metabolism^13^. In humans, there are three common alleles of *APOE*: ε2, ε3, and ε4. The ε4 allele is carried by nearly 20% of the population, and confers up to 15× increase in risk for AD compared to E3 homozygotes^14^.

Amyloid plaques trigger transcriptional changes in nearby microglia, inducing a shift toward a pathological signature^15,16^. Similar neurodegenerative profiles have been described across several independent studies, being termed Activated Response Microglia (ARM)^15^, neurodegenerative microglia (MGnD)^17^, or Disease Associated Microglia (DAM)^18^. Interestingly, many of the genes within the aforementioned profiles belong to metabolic pathways, including core genes such as *Ch25h, Fabp5, Hexb, Lpl*, and *Apoe* itself^19–24^. While several studies attempted to translate these findings to postmortem human brain tissue, they found very little overlap between human AD microglial gene signatures and those identified in mouse models^25–27^. A glaring exception to this lack of overlap was *Apoe/APOE*, whose expression is amplified in neurodegenerative conditions across all studies and species – indicating that it is a universal, core transcriptomic ‘switch’ within AD-associated microglia. However, it remains unclear whether isoform-specific differences in this process underlie the harmful effect of E4 in AD.

A handful of previous studies have examined the role of human *APOE* on the mouse brain transcriptome and metabolome^28–32^, and a recent study inferred strong, glial-driven *APOE* genotype effects from whole-tissue bulk RNA sequencing of postmortem human brains^33^. Together these important studies highlight both amyloid-dependent and independent roles for *APOE*, age, and brain region in metabolic and immune changes. However, their reliance on pre-selected brain regions and bulk homogenates limits insight into specific glial cell type contributions and lacks sub-regional anatomic resolution.

Here, we employed a single-source experimental design that combines bulk, single-cell, and spatial transcriptomics with cell-specific and spatially resolved metabolic analyses in order to systematically address the role of *APOE* across age, neuroinflammation, and AD pathology. Both bulk tissue and single-cell RNA sequencing (scRNAseq) highlighted immunometabolic changes across the *APOE4* glial transcriptome. Although aged E4 mice lack any observable AD pathology, we note that the gene signature expressed by their microglia: i) includes a robust increase in *APOE*, ii) is overrepresented by genes involved in glucose metabolism, lipid processing and innate immunity, and iii) substantially overlaps with gene signatures previously described in both mouse models and human AD microglia. Further, exposing mice to a systemic inflammatory challenge resulted in a metabolically distinct response within E4 microglia. Using metabolomics and functional metabolic assays, we show that E4 microglia are inherently pro-glycolytic and *HIF1α*-high, displaying a metabolic profile classically associated with activated (M1/M[LPS, IFNγ, TNFα]) myeloid cells. We then crossed E3 and E4 mice to amyloid overexpressing 5XFAD mice and utilized spatial transcriptomics (ST) to determine anatomically salient changes in gene expression. ST highlighted the cortex and hippocampus as particularly sensitive to *APOE4* and revealed a unique response to amyloid in the E4 brain characterized by microglial activation and widespread alterations in lipid metabolism. Matrix-assisted laser desorption ionization (MALDI) mass spectrometry imaging (MSI) confirmed age-, *APOE*-, and region-specific alterations in lipid metabolism, particularly in multiple phospholipid species. Finally, we provide researchers with an interactive web-based resource (www.LJohnsonLab.com/database) in which to explore the effects of *APOE* across aging, neuroinflammation, and AD pathology via bulk, single-cell, and spatial transcriptomic datasets. Together, our findings link two phenomena consistently tied to AD (metabolic dysfunction and neuroinflammation) to the strongest genetic predictor of late onset AD (E4), emphasizing a role for *APOE* in regulating glial immunometabolism.

## Results

### *APOE4* drives immunometabolic changes across the glial transcriptome

In order to systematically examine the effect of *APOE* genotype across aging, neuroinflammation, and AD pathology, we designed a single-source multi-omics approach that combined bulk and scRNAseq with cell-specific metabolomics and serial-section ST, histopathology, and MALDI MSI (**Fig. 1a**). To examine the effects of *APOE* across the lifespan, we began with bulk sequencing of whole brain tissue homogenates from young (3 mo.), middle-aged (12 mo.), and aged (24 mo.) mice expressing human E3 or E4. We identified a few hundred differentially expressed genes (DEGs) between E4 and E3 brains (**Fig. 1b-c**), including previously reported genes such as *Serpina3n* (**Fig. 1d**)^28,30^. To better understand these gene expression changes at a systems-level, we performed a pathway analysis of E4 vs E3 DEGs. Nine out of the top 10 KEGG terms fell under the umbrella pathways of “metabolism” or “immune system” (**Fig. 1e**).

**Figure 1.**
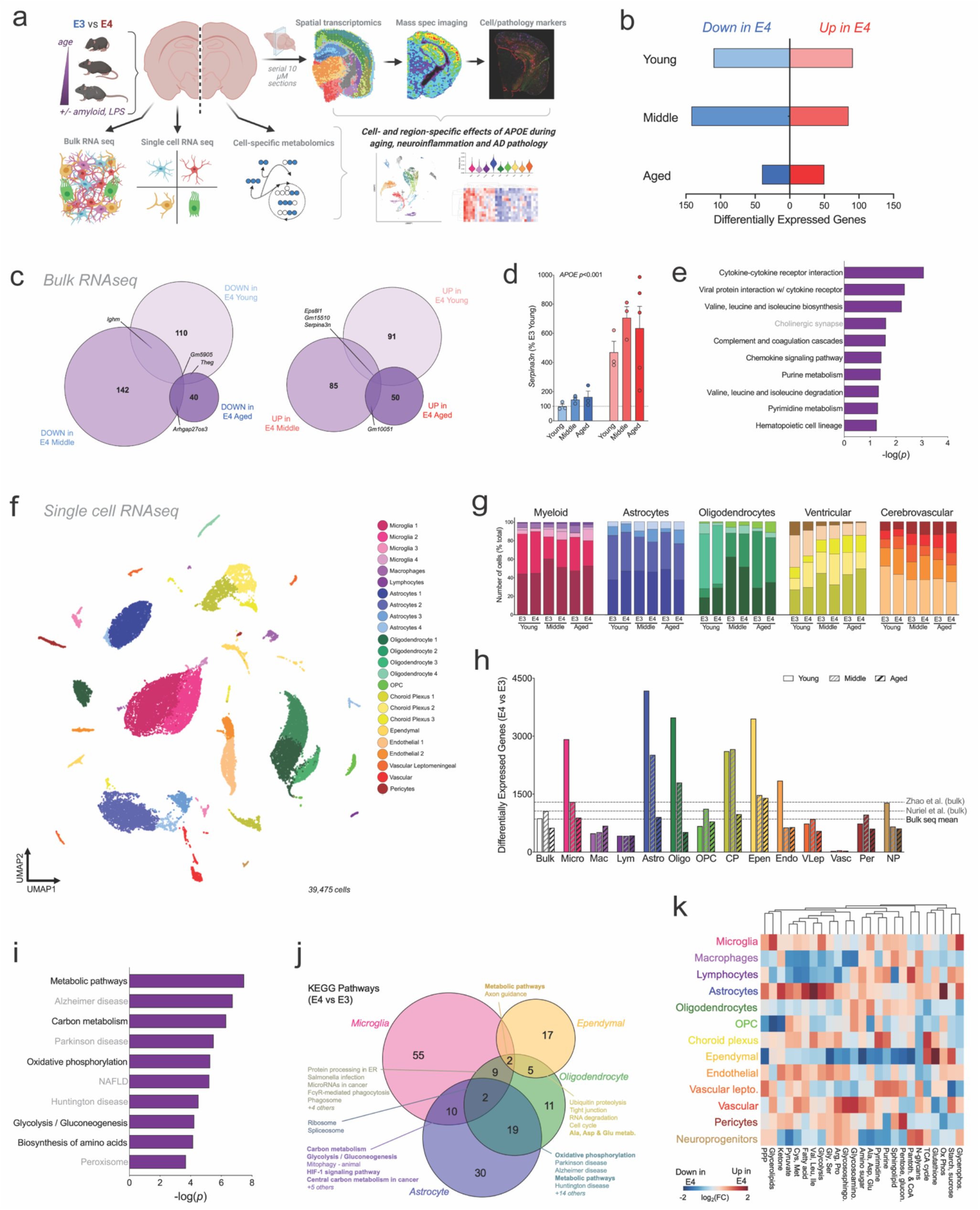
APOE4 drives immunometabolic changes across the glial transcriptome. a) Experimental design. Brains from APOE3 and APOE4 mice were analyzed across the lifespan (3, 12 and 24 months of age), and in the presence of an inflammatory challenge (LPS) or AD pathology (amyloid overexpression). b-c) Number (b) and overlap (c) of differentially expressed genes (DEG) between E3 and E4 brains at each age (bulk RNAseq). Each circle is a comparison in young (light purple), middle aged (purple), or aged (dark purple); relative size corresponds to total DEGs. d) Gene expression of *Serpina3n* in whole brain. e) The top 10 KEGG pathways most significantly altered by *APOE4* in whole brain tissue. Terms in bold fall under KEGG umbrella pathways of “metabolism” or “immune system”. f) UMAP showing 24 clusters classified based on canonical gene expression markers. g) Number of cells per cluster. h) DEGs between E3 and E4 brains within each cell type at each age (scRNAseq). Young, open bars; Middle age, grey dashed bars; Aged, black dashed bars. Dashed lines indicate the number of DEGs in this, and two previous, bulk seq analyses^28–29^. i) The top 10 KEGG pathways most significantly altered by *APOE4* across all glia (scRNAseq). j) Venn diagram showing overlap of KEGG pathways differentially expressed between E4 and E3 in the four cell types most affected by *APOE4*. Numbers represent number of significantly altered pathways in each cell type. The top 5 overlapping KEGG pathways are listed for each intersection. k) Heatmap of the top 25 metabolic pathways altered by *APOE* in each cell type. Pathways in red show increased expression in E4 cells; blue, decreased.

To identify cell-specific contributions to these whole tissue gene expression changes, we performed scRNAseq on the same brains analyzed for bulk sequencing. Dimensionality reduction using uniform manifold approximation and projection (UMAP) identified 24 clusters which were assigned to one of 13 unique cell types using established gene markers (**Fig. 1f; Extended Data Fig. 1a**). While age-related enrichment of some clusters was observed, cell numbers were similarly distributed across *APOE* genotypes (**Fig. 1g; Extended Data Fig. 1b-c**). Analysis of DEGs highlighted astrocytes, oligodendrocytes, ventricular cells, and microglia as the cell types most affected by *APOE* (**Fig. 1h**). Similar to the bulk tissue, the number of DEGs decreased with age across several cell types (i.e., the effects of E4 were more pronounced in younger brains). A pathway analysis of DEGs across all cells together once more highlighted metabolism, particularly the central carbon metabolic pathways of oxidative phosphorylation (OxPhos) and glycolysis (**Fig. 1i**). Further, many of these differentially expressed metabolic pathways and hypoxia inducible factor 1 (HIF-1) signaling were shared across multiple cell types (**Fig. 1j; Table 1**).

Calculation of metabolic pathway activity for each individual cell using AUCell^34^ revealed the effect of E4 to be highly variable and cell-specific. For example, astrocytes showed more pronounced E4-associated
increases in branched chain amino acid metabolism and OxPhos, while microglia displayed robust E4-associated increases in glycerolipid metabolism and glycolysis (**Fig. 1k**). Together, these results suggest that the major transcriptomic changes driven by *APOE4* involve glial metabolism and the immune response.

### *APOE* expression is selectively upregulated in aged E4 microglia

We next asked if expression of *APOE* itself varied across the lifespan in E3 and E4 glia. While *APOE* expression did not vary by age or genotype in whole brain tissue (**Fig. 2a**), several cell-specific changes were noted. First, while *APOE* was predominantly expressed by astrocytes, many other cell types showed measurable levels of both *APOE* (**Fig. 2b-d**) and its respective binding partners (**Fig. 2e-f; Extended Data Fig. 2a-b**). Strikingly, while most cell types had subtle, if any, changes in *APOE* expression across the lifespan, aged E4 microglia showed a unique and dramatic upregulation *APOE* (**Fig. 2g-h**). Re-clustering of astrocytes and microglia showed that this upregulation was limited to a distinct sub-population of microglia (Mi_6) (**Fig. 2i-k; Extended Data Fig. 2c**). Finally, the *APOE* signaling network in Mi_6, along with almost all astrocyte clusters, was markedly upregulated in both outgoing (*APOE*) and incoming (*Ldlr, Lrp1*, etc.) signal strength in the aged E4 brain (**Extended Data Fig. 2d**). In sum, these findings highlight an age-related increase in *APOE* expression – in the absence of AD pathology – that is unique to E4 microglia.

**Figure 2.**
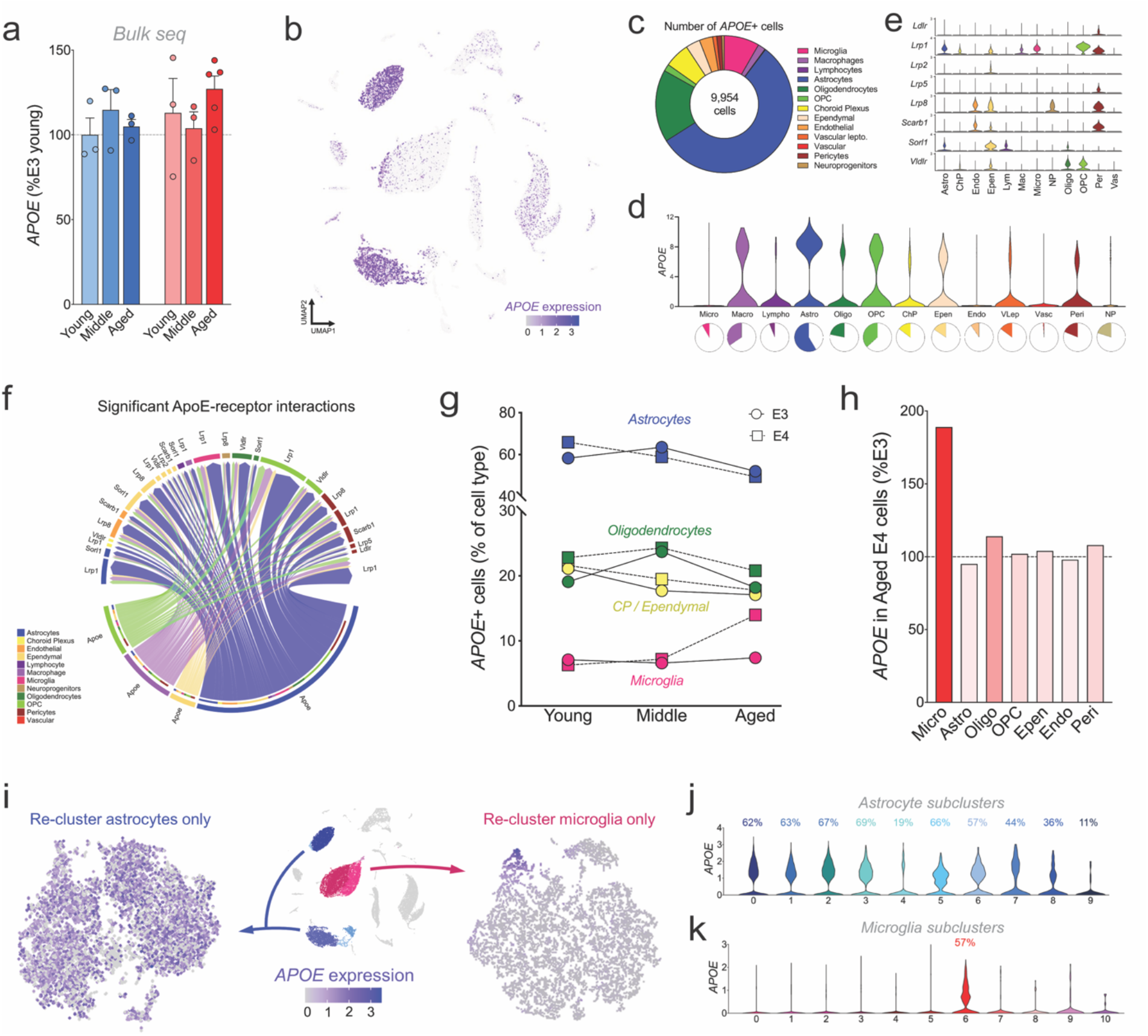
*APOE* expression is selectively upregulated in aged E4 microglia. a) *APOE* expression in whole brain tissue across the lifespan (‘Bulk seq’). *p*=0.39 for *APOE, p*=0.68 for age. b) UMAP showing *APOE* expression across all cell types. c-e) Donut chart of all 9,954 cells expressing *APOE* at a detectable level (“*APOE*+”). Color slices denote the fraction of total *APOE*+ cells belonging to each cell type. d) Violin plots showing *APOE* expression across each cell type. Pie charts (bottom) show the percent of each cell type that are *APOE*+. e) Expression of established ApoE receptors within each glial cell type. f) Circos plot showing significant ApoE-receptor interactions as calculated by CellChat. Astrocytes (bottom, blue) express the majority of *APOE* and signal to a variety of ApoE receptors across multiple cell types (top). Bottom: outer circle color denotes the cell type expressing *APOE*, while the inner circle color denotes the cell type expressing the corresponding receptor (ex. *Ldlr*). g) *APOE* expression ‘over time’ in the primary *APOE* expressing cell types (from c). The percent of *APOE*+ cells within each glial cell type is shown in young, middle aged, and aged E3 and E4 brains. h) *APOE* expression in aged E4 glia as a percent of E3 cells. Expression of *APOE* is selectively upregulated in aged E4 microglia. i-k) *APOE* expression is restricted to a subset of microglia. i) tSNE plots of astrocytes (left) and microglia (right), showing expression of *APOE* within each cell type. *APOE* is highly expressed across the majority of astrocyte sub-clusters (j), while expression in microglia (k) is restricted to Mi_ 6.

### *Hif1α*-high, ‘DAM-like’ microglia are increased in the aged E4 brain

To distinguish microglial genes that significantly change with age and/or E4, we calculated gene scores for each individual microglia. Strikingly, genes that are upregulated in both advanced age and E4 were heavily enriched for DAM/MGnD genes (**Fig. 3a**). Further, E4-specific changes in the microglia transcriptome substantially overlapped with multiple AD-relevant gene lists from both mouse and human studies (**Fig 3b; Table 1**). Interestingly, we observed a ‘flip’ in expression patterns for many AD-associated genes whereby young E4 microglia had higher expression but aged E4 microglia had lower expression compared to E3, or vice versa (**Fig. 3c**). Due to its unique upregulation of *APOE*, we next focused on microglia cluster 6 (Mi_6). Remarkably, the biomarkers that defined Mi_6 were almost exclusively genes associated with the DAM/MGnD signature, including metabolic genes *Lpl, Ch25h, Fabp5* and *APOE* itself (**Fig. 3a; Table 1**). Mi_6 was highly enriched in aged E4 mice relative to E3 (**Fig. 3b; Extended Data Fig. 3a-b**) and a pathway analysis of the Mi_6 biomarkers highlighted “Alzheimer’s disease” and metabolic pathways including “cholesterol metabolism” and “HIF1 signaling” (**Fig. 3f**).

**Figure 3.**
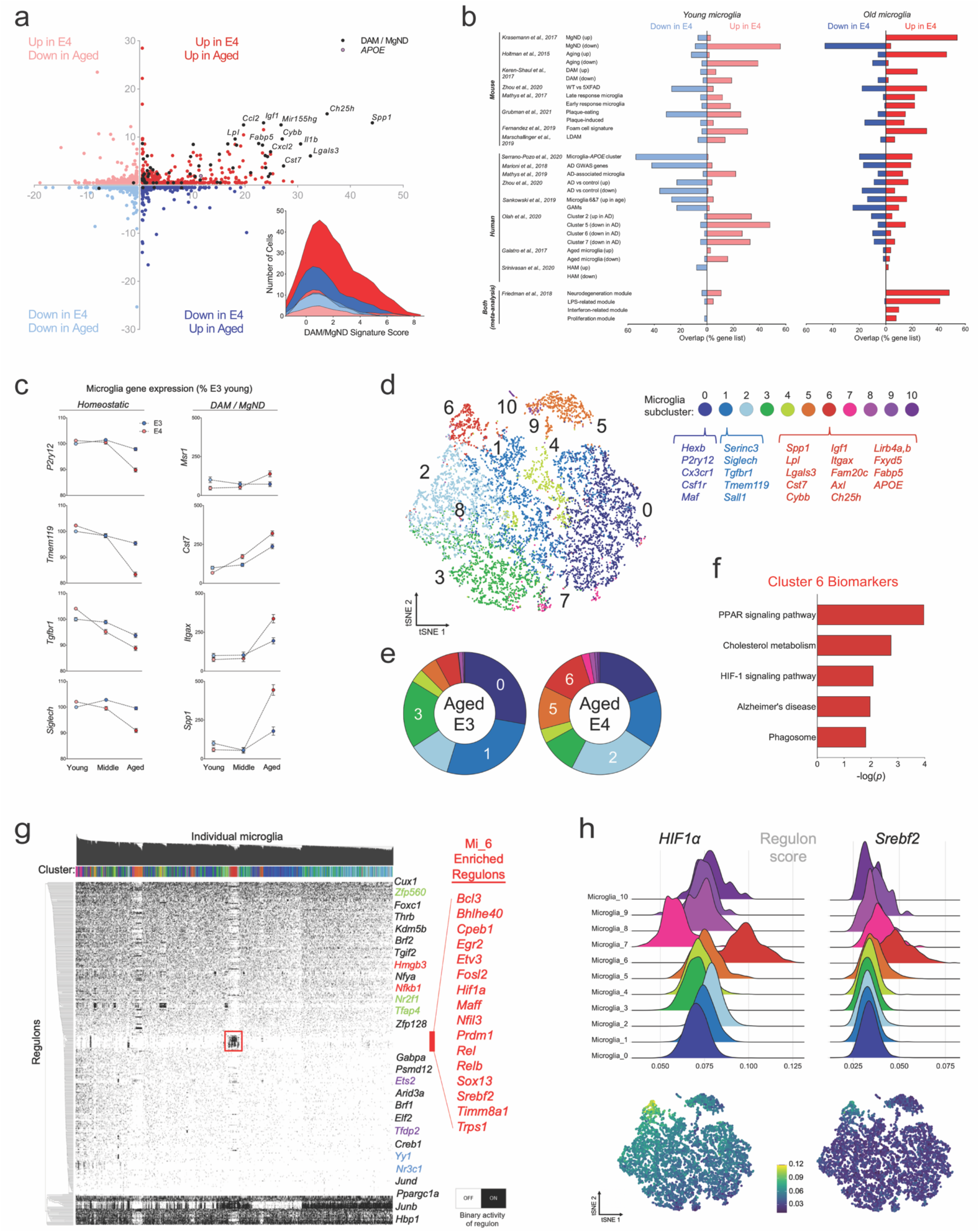
Age and *APOE4* are associated with an increase in ‘DAM-like’ microglia. a) Gene score plot showing DEGs between E4 vs E3 microglia (y-axis) and aged vs young microglia (x-axis). Genes labeled in black are common to both DAM/MgND phenotypes. Inset: Ridge plot showing DAM/MgND score for each individual microglia as calculated by AUCell. b-c) E4-specific changes in the microglia transcriptome substantially overlap with AD-relevant gene lists from mouse and human studies. b) Overlap of published gene lists with DEGs (E4 vs E3) in young (left) and aged (right) microglia. c) Expression of select ‘homeostatic’ and DAM/MgND genes in young, middle and aged microglia. d-f) Aged E4 microglia are enriched for a sub-cluster of cells with a DAM-like expression profile (Cluster 6; Mi_6). d) tSNE of microglia sub-clusters. Top biomarkers for the ‘homeostatic’ clusters (0,1) and the ‘DAM-like’ cluster 6 are displayed beneath the cluster labels. e) Donut charts showing the distribution of aged E3 (left) and aged E4 (right) microglia within each sub-cluster. Clusters labeled in white are enriched in the respective group. f) Top 5 gene ontology (GO) terms associated with the biomarkers that define Mi_6. g) SCENIC was used to reconstruct active regulons in each individual microglia and meaningfully cluster cells based on shared activity patterns (binarized). Mi_6 is defined by selective high activity of 16 TFs (red box; ‘Mi_6 Enriched Regulons’) and the relative absence of activity of other TFs. h) Ridge plots (top) or tSNE (bottom) showing regulon activity scores for *HIF1α* (left) and *Srebf2* (right).

In order to identify potential upstream regulators that define these various microglia clusters, we used SCENIC to reconstruct active regulons (i.e., transcription factors (TFs) and their target genes) in individual microglia^34^. SCENIC revealed a clear and distinctive clustering of Mi_6 defined by 16 regulons (**Fig. 3g-h**). Intriguingly, several of these regulons have been previously implicated in AD (*Bhlhe40*)^17^, regulate metabolic pathways (*Timm8a1*^35^, *Srebf2* ^36^), or both (*Hif1α*^37^) (**Fig. 3 g-h**). *HIF1α* in particular was substantially upregulated in Mi_6 (**Fig. 3h**) and was positively correlated with the cell’s DAM/MGnD score (**Extended Data Fig. 3c**). Given the central role of these regulons in metabolism, we next sought to characterize metabolic activity within each cluster. A heatmap of metabolic pathway scores revealed Mi_6 as the cluster with the highest expression of central carbon pathways, including glycolysis (**Extended Data Fig. 3d-e**). Together, these data show that even in the absence of overt AD pathology, age, and E4 are sufficient to drive changes in microglia that i) overlap with both mouse and human AD-relevant gene lists, ii) strongly resemble a DAM phenotype, and iii) prominently feature distinct shifts in the regulation of glucose and lipid metabolism.

### E4 microglia have a distinct metabolic response to an inflammatory challenge

Given their unique metabolic transcription profile and DAM-like signature, we next asked whether E4-expressing microglia would differentially respond to an inflammatory challenge. Twenty-four hours following a peripheral injection of LPS or saline, we harvested brains from E3 and E4 mice and performed scRNAseq on microglial populations (**Fig. 4a**). Microglia from the E4 LPS brains showed a remarkably distinct metabolic profile, with increased activity across multiple pathways of amino acid, sugar, and fatty acid metabolism (**Fig. 4b**). At the subpopulation level, treatment with LPS resulted in several distinct clusters of microglia, including two clusters enriched in E3 LPS brains (5,7) and two found almost exclusively in E4 LPS brains (8,11) (**Fig. 4c-d**). Notably, the DEGs defining these E4 LPS-enriched clusters correspond to gene ontology terms related to mitochondrial function, aerobic respiration, and energy production (**Fig. 4e**). These E4 LPS-enriched clusters also showed high expression of genes belonging to OxPhos and glycolysis pathways (**Fig. 4f**). In total, these data suggest that compared to E3, expression of E4 leads to a robust and distinct metabolic response by microglia to an inflammatory challenge.

**Figure 4.**
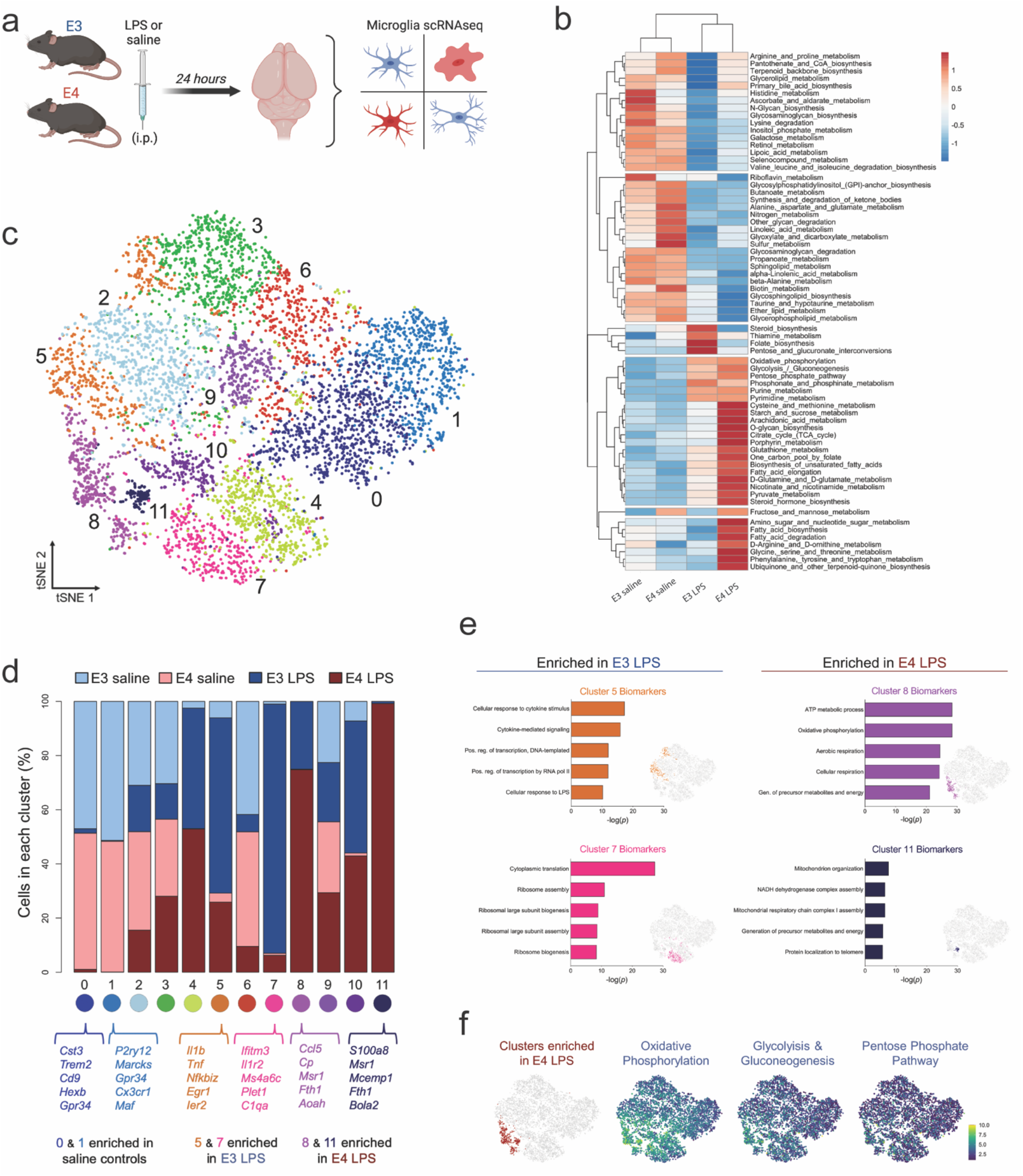
APOE4 microglia are metabolically distinct in response to an inflammatory challenge. a) Experimental design. E3 and E4 mice were injected with lipopolysaccharide (LPS; 5mg/kg) or saline, and brains dissected 24 hours later for scRNAseq. b) Heatmap showing expression of KEGG metabolic pathways in microglia from LPS or saline treated mice. c-d) E3 and E4 brains show enrichment of distinct microglia sub-clusters following LPS treatment. B) tSNE plot of microglia from LPS or saline treated E3 and E4 mice. Colors highlight the 12 identified microglia sub-clusters. c) Stacked bar plot showing distribution of experimental groups within each microglia sub-cluster. Top 5 biomarkers for the two ‘homeostatic’ (0,1), E3-enriched (5,7), and E4-enriched (8,11) clusters are listed below. e-f) E4 LPS microglia are associated with energy production and oxidative phosphorylation pathways. e) Top 5 gene ontology terms associated with the two E3 LPS-enriched (left; 5,7) and two E4 LPS-enriched (right; 8,11) clusters. f) tSNE plots showing higher expression of central carbon (i.e. energy production) pathways (right) in sub-clusters enriched in the E4 LPS brain (left).

### E4 microglia have increased aerobic glycolysis and higher *HIF1α* expression

We next sought to determine if these differences in gene expression would be functionally reflected in altered metabolism between E4 and E3 primary microglia (**Fig. 5a**). Using a targeted metabolomics approach, we identified five metabolites that were significantly upregulated in E4 microglia (lactate, succinate, glutamine, tyrosine, and threonate) and one significantly downregulated (itaconate) (**Fig. 5b-c; Extended Data Fig. 4a**).

**Figure 5.**
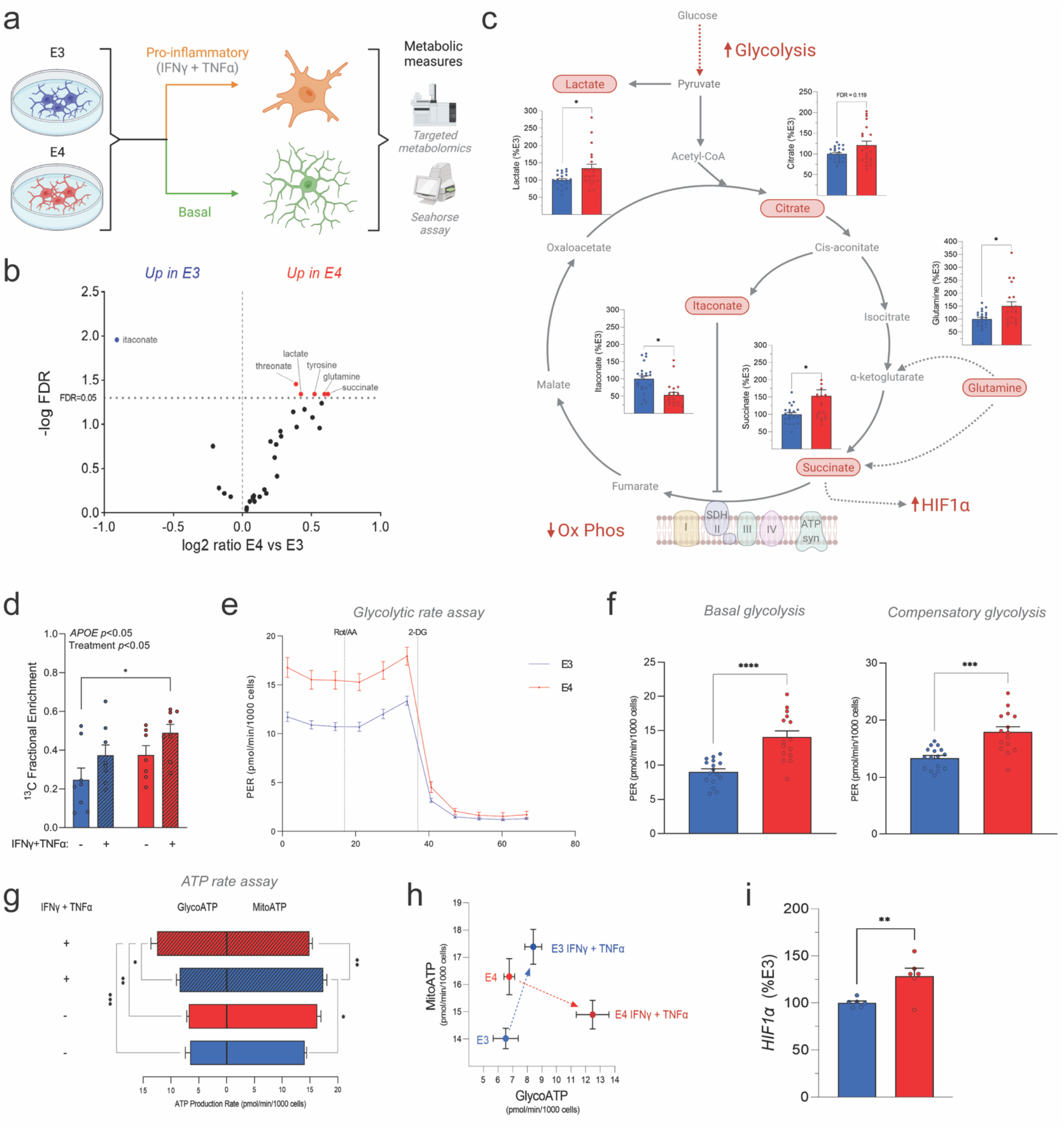
E4 microglia have increased aerobic glycolysis and higher *Hif1α* expression. a) Experimental design. Primary microglia were isolated from E3 and E4 mice and stimulated *in vitro* with a pro-inflammatory (20ng/ml IFNγ + 50ng/ml TNFα) cytokine cocktail prior to Seahorse analysis or targeted metabolomics (both steady-state and stable-isotope resolved metabolomics). b-c) Targeted metabolomics on E3 and E4 microglia (n=21-22). b) Volcano plot showing changes in steady-state metabolites. c) Schematic of TCA cycle and glycolysis. Pathways and metabolites associated with pro-inflammatory immunometabolism are highlighted in red, with corresponding bar graphs for E3 and E4 steady-state metabolites overlaid on each. d) Stable-isotope tracing reveals increased fractional enrichment of fully-labeled (m+3) lactate in pro-inflammatory treated E4 microglia (n=7-8) after 2 hours. e-f) Proton efflux rate (PER) (pmol/min/1000 cells), a measure of glycolysis, measured over time in E3 and E4 microglia during the Glycolytic Rate Assay (Agilent). f) E4 microglia showed higher basal glycolysis (left) and compensatory glycolysis (right) compared to E3 controls (n=15-16). g-h) ATP production rate (pmol/min/1000 cells) measured during the ATP Rate Assay (Agilent) in E3 and E4 microglia, with glycolytic ATP production (GlycoATP) to the left of the y-axis and mitochondrial ATP production (MitoATP) to the right (n=5-9). h) XY plot with MitoATP displayed on y-axis and GlycoATP displayed on X-axis. E4 microglia respond to stimulus by dramatically increasing GlycoATP and decreasing MitoATP (red dashed arrow) whereas E3 microglia respond with only a slight increase to GlycoATP and instead show a dramatic increase in MitoATP (blue dashed arrow). i) Quantitative RT-PCR analysis shows increased *Hif1α* gene expression in E4 primary microglia (n=6). *p<0.05, **p<0.01, ***p<0.001, ****p<0.0001. 2-way ANOVA (d,g), two-tailed T-test (f,i), or two-tailed T-test adjusted for multiple comparisons (indicated as FDR, False Discovery Rate) (b-c).

Of note, lactate accumulates in cells undergoing increased aerobic glycolysis, such as pro-inflammatory activated macrophages^11,12^. Citrate and succinate also accumulate in pro-inflammatory macrophages due to a break in the TCA cycle^11,12^. Succinate was significantly increased in E4 microglia, while concentrations of itaconate, which activates downstream anti-inflammatory and antioxidant signaling pathways^38–40^, was lower (**Fig. 5b-c**).

To ascertain whether these differences in steady-state metabolite pool sizes were part of a more dynamic alteration in metabolic flux within E4 microglia, we next turned to stable-isotope resolved metabolomics. E3 and E4 microglia were stimulated with a combination of interferon-γ (IFNγ) and tumor necrosis factor α (TNFα) in the presence of ^13^C-glucose and incorporation of ^13^C in downstream metabolites was measured. This revealed a significant increase in fully labeled (m+3) ^13^C-lactate due to both *APOE4* and pro-inflammatory treatment, indicating increased flux of glucose through aerobic glycolysis (**Fig. 5d**).

To functionally assess the effect of *APOE* on microglial metabolism, we employed the Seahorse platform to measure glycolysis, mitochondrial respiration, and the relative contribution of each pathway to ATP production. Interestingly, we noted that E4 microglia showed higher rates of basal and compensatory glycolysis compared to E3 (**Fig. 5e-f**) and had lower maximal respiration and spare respiratory capacity (**Extended Data Fig. 4c-f**). Additionally, E4 microglia responded to a pro-inflammatory stimulus by dramatically increasing glycolytic ATP production at the expense of decreased mitochondrial production. In contrast, E3 microglia significantly increased mitochondrial ATP production following stimulation (**Fig. 5g-h**). These data suggest that E4 microglia rely exclusively on a substantial upregulation of glycolysis to support the increased energy demand of the pro-inflammatory response, whereas E3 microglia demonstrate increased metabolic flexibility (**Fig. 5g-h**).

Finally, the increased succinate and clear functional shift toward aerobic glycolysis in E4 microglia is congruent with the increased activity of the *Hif1α* regulon in the E4 microglia SCENIC data. When stabilized by a pro-inflammatory stimulus (and/or succinate), the HIF1α transcription factor complex translocates to the nucleus and activates many genes important for increasing glycolysis^41^. In agreement with this pro-glycolytic E4 phenotype, quantitative RT-PCR revealed increased expression of *Hif1α* in E4 microglia compared to E3 (**Fig. 5i**). Together these data highlight functional metabolic reprogramming whereby E4 microglia are inherently pro-glycolytic and anti-oxidative, a phenotype that mirrors classically activated macrophages.

### Spatial transcriptomics identifies unique cortical and hippocampal signatures of *APOE4*, age, and amyloid overexpression

We next leveraged Visium ST technology to assess gene expression across coronal brain sections from young, aged, and 5XFAD amyloid overexpressing E3 or E4 mice (**Fig. 6a; Extended Data Fig. 5a**). A total of 16,979 spots were analyzed across six brains, and high dimensionality reduction identified 18 total clusters. This included 17 anatomically conserved clusters that expressed canonical region-specific markers and mapped exceptionally well to the Allen Brain Atlas (**Fig. 6a-b; Extended Data Fig. 5b-c**). Intriguingly, the final cluster (Cluster 11) was found almost exclusively in the E4 5XFAD brain and was primarily localized within cortical regions (**Fig. 6c**). This cluster, which we term a ‘disease associated’ signature, was defined by biomarkers enriched in pathways related to lipid metabolism, synapse pruning, neuronal death, and microglial activation (**Fig. 6d**). When we mapped these cluster 11 biomarkers back to our scRNAseq data, the signature was exclusively and highly expressed by microglia, with the Mi_6 cluster showing peak expression (**Fig. 6c**).

**Figure 6.**
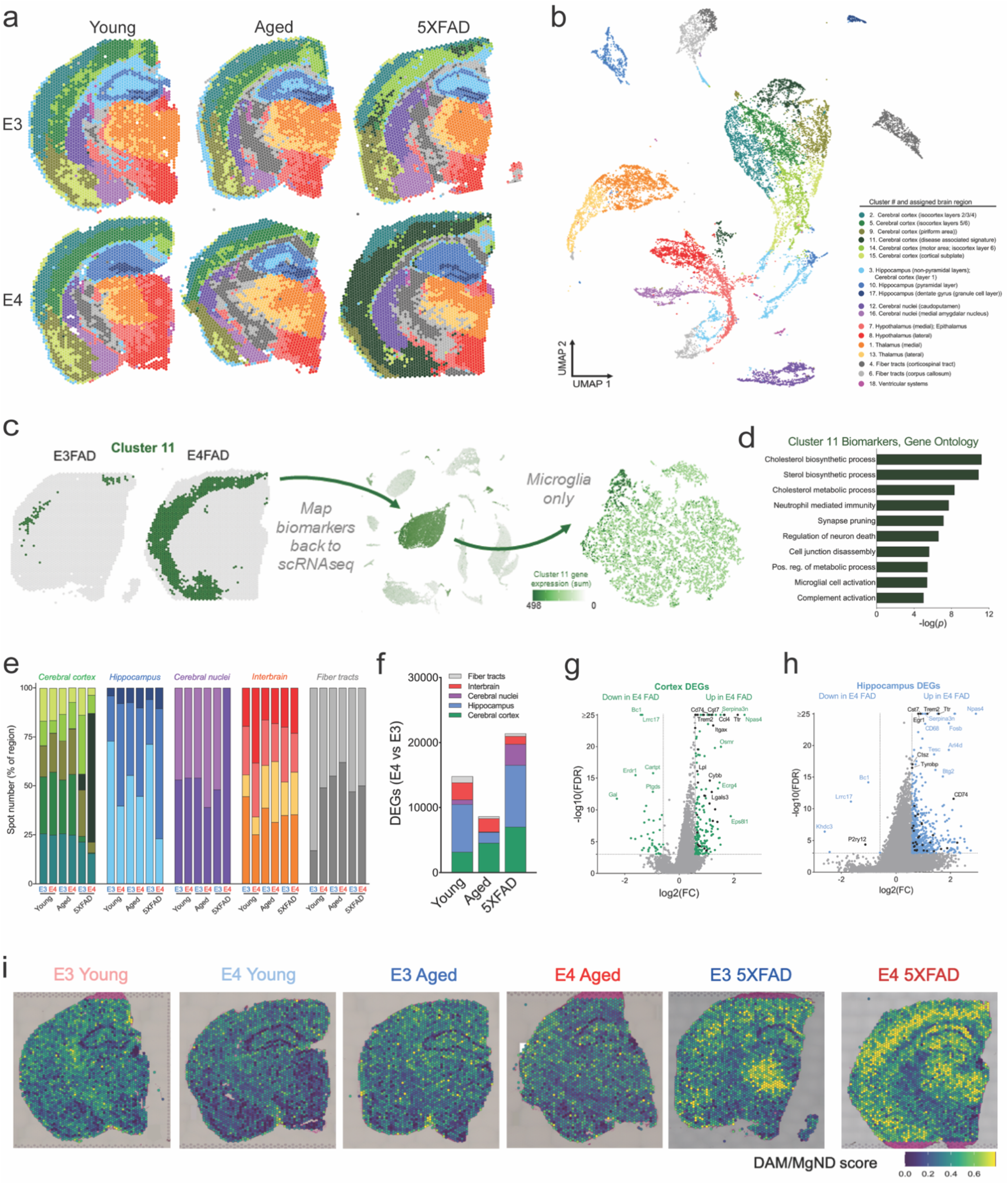
Spatial transcriptomics highlights unique cortical and hippocampal signatures of *APOE4*, age and amyloid overexpression. a-b) Spatial transcriptomics (ST) identifies 17 unique clusters that are anatomically conserved, plus one unique cortical cluster primarily restricted to E4FAD mice (Cluster 11, dark green). a) Spatial transcriptomic plots of brain sections from young, aged, and amyloid overexpressing E3 and E4 mice. b) UMAP plot of all 79,980 spots analyzed across all 6 brains. Clusters were assigned labels based on anatomical concurrence to the Allen Brain Atlas. c-d) Cluster 11 is enriched in the E4FAD brain and consists of genes related to lipid metabolism and microglial activation. c) E3FAD and E4FAD brains showing spots belonging to Cluster 11. Cluster 11 biomarker genes were re-plotted to scRNAseq data, showing highest expression in microglia, specifically in Mi_6. d) Top 10 gene ontology terms for Cluster 11, highlighting pathways of lipid metabolism and immune activation. e) Number of spots within each cluster for each experimental group. Clusters are organized by respective brain regions. f-h) E4 drives gene expression changes primarily in the cortex and hippocampus. f) Differentially expressed genes (DEGs) between E4 and E3 brains within each brain region. g-h) DEGs within the cortex (g) and hippocampus (h) of the 5XFAD mice. Genes labeled in black correspond to DAM/MgND genes. i) ST plots showing DAM/MgND scores for each spot (calculated with AUCell).

We next assigned spots to one of five primary brain regions, noting that the majority of E4 vs E3 DEGs were found in the cerebral cortex and hippocampus (**Fig. 6e-f; Extended Data Fig. 6a-c**). Both regions featured a robust upregulation of genes in the E4 5XFAD compared to E3 5XFAD brain, many of which were DAM/MGnD genes (**Fig. 6g-i**). Additionally, gene markers of glial reactivity previously linked to *APOE4* were similarly upregulated in E4 aged and E4 5XFAD brains (**Extended Data Fig. 6d**). Taken together, these results i) support an E4-associated increase in microglial activation, ii) highlight cortical and hippocampal regions as areas most affected by *APOE*, and iii) reveal a unique E4 response to amyloid pathology.

### *APOE4* exacerbates plaque-induced microglial activation and lipid metabolism

To determine whether this unique E4 5XFAD transcriptional profile was spatially linked to AD pathology, we stained for amyloid plaques across the 10μM section immediately adjacent to that subjected to ST and assigned each spot a numerical plaque intensity score (**Fig. 7a**). A correlation analysis revealed a number of genes that either positively or negatively tracked with plaque intensity. Notably, in the E4 brain, this included strong positive correlations with markers of glial reactivity and a 3-fold increase in the number of significantly correlated DAM genes (**Fig. 7b**). A pathway analysis of significantly correlated genes showed numerous terms shared by both E3 and E4 (purple), as well as some unique to E4 (red) or E3 (blue) (**Fig. 7c**). Shared pathways included terms related to synaptic transmission (negatively correlated) or synapse pruning and microglial activation (positively correlated) (**Fig. 7c**). Interestingly, pathways unique to E4 were predominantly related to lipid metabolism (**Fig. 7c**).

**Figure 7.**
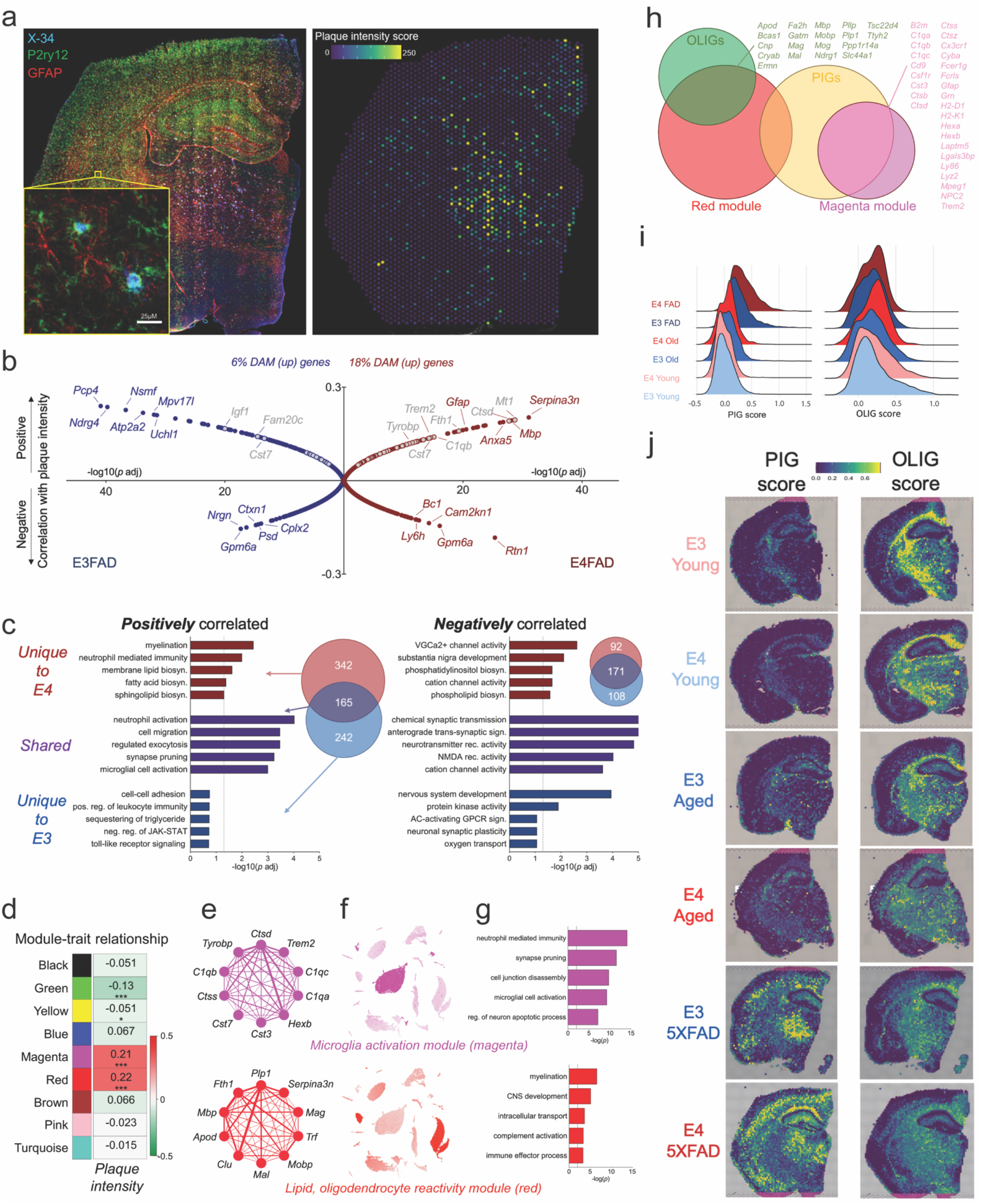
*APOE4* exacerbates plaque-induced microglial activation and alterations in lipid metabolism. a) E4FAD brain stained with P2ry12 (green; microglia), GFAP (red; astrocytes), and X-34 (blue) to demarcate amyloid plaques (left). X-34 intensity was quantified to generate a ‘plaque intensity score’ for each individual spatial transcriptomic spot (right). b) Gene correlation with plaque intensity in E3FAD (blue, left) and E4FAD (red, right) brains. Y-axis values represent correlation coefficients, with genes at the top of the graph positively correlated with plaque intensity, and genes at the bottom negatively correlated. Distance from center on the x-axis represents significance of the correlation (-log10(*p* adjusted)). DAM/MgND genes are noted in gray. c) Top 5 gene ontology terms for genes that were positively (left) or negatively (right) correlated with plaque intensity. Some GO terms were uniquely correlated with E4 (red), some uniquely correlated with E3 (blue), and some correlated with plaque intensity regardless of *APOE* genotype (purple). Venn diagrams show overlap between genes correlated with plaque intensity in E4FAD (red circles) or E3FAD (blue circles) brains. d-g) Gene networks associated with plaque intensity. d) The correlation between module eigengenes (MEs) and amyloid plaque intensity. Values in the heatmap are Pearson’s correlation coefficients, and stars represent significant correlations: *p < 0.05; ***p < 0.001. Modules with positive values (red) indicate positive correlation of MEs with plaque intensity, modules with negative values (green) represent a negative correlation. E) Network plots of the top 10 genes with the highest intramodular connectivity (hub genes) in the magenta (top) and red (bottom) modules. f) UMAP plots map expression of module gene lists (sum) back to the scRNAseq dataset. g) Top 5 gene ontology terms associated with the magenta or red modules. h) Venn diagrams showing overlap of red and magenta modules with oligodendrocyte (OLIG) and plaque induced gene (PIG) lists from [REF]. Overlapping genes are listed. i-j) The E4FAD brain has a higher PIG score and lowest OLIG score. i) Ridge plots showing PIG (left) and OLIG (right) scores for each experimental group. j) Spatial expression of PIG (left) and OLIG (right) gene lists.

A weighted gene co-expression network analysis (WGCNA) highlighted two networks (green, yellow) containing genes related to ion channels and synaptic transmission that were negatively associated with plaque intensity, and two networks (magenta, red) that were positively associated (**Fig. 7d-g; Extended Data Fig. 7a-b**). The magenta network we termed a ‘microglia activation module’ as it mapped almost exclusively to microglia in our scRNAseq database, was enriched for DAM genes, and was associated with gene ontology terms related to synapse pruning, neuron apoptosis and microglia activation (**Fig. 7e-g**, *top*). In contrast, the red ‘lipid, oligodendrocyte reactivity’ module mapped predominantly to oligodendrocytes, and included markers of lipoprotein transport, myelin, and glial reactivity (**Fig. 7e-g**, *bottom*).

Both the red and magenta modules were more highly expressed across the E4 5XFAD brain relative to E3 (**Extended Data Fig. 7a**) and intriguingly they substantially overlapped with the oligodendrocyte (OLIG) and plaque induced gene (PIG) networks identified from a previous ST study of AD mouse and human brains^16^ (**Fig. 7h**). Interestingly, the PIG score was lowest in the young brains, increased slightly with age, and was highest in the E4 5XFAD brain; with the OLIG scoring following the opposite trend (**Fig. 7i-j**). In sum, these data highlight a unique E4 response to increasing amyloid pathology characterized by increased microglial activation and alterations in lipoprotein and lipid metabolism.

### MALDI MSI confirms *APOE*-, age-, and amyloid-associated changes in lipid metabolism

In whole brain tissue, we noted significant effects of *APOE*, age, and their interaction on the expression of multiple lipid metabolism pathways (**Fig. 8a**). At the single-cell level, changes in lipid metabolism were most pronounced in microglia, specifically in glycerophospholipids, with aged E4 microglia having the highest expression (**Fig. 8b; Extended Data Fig. 8a**).

**Figure 8.**
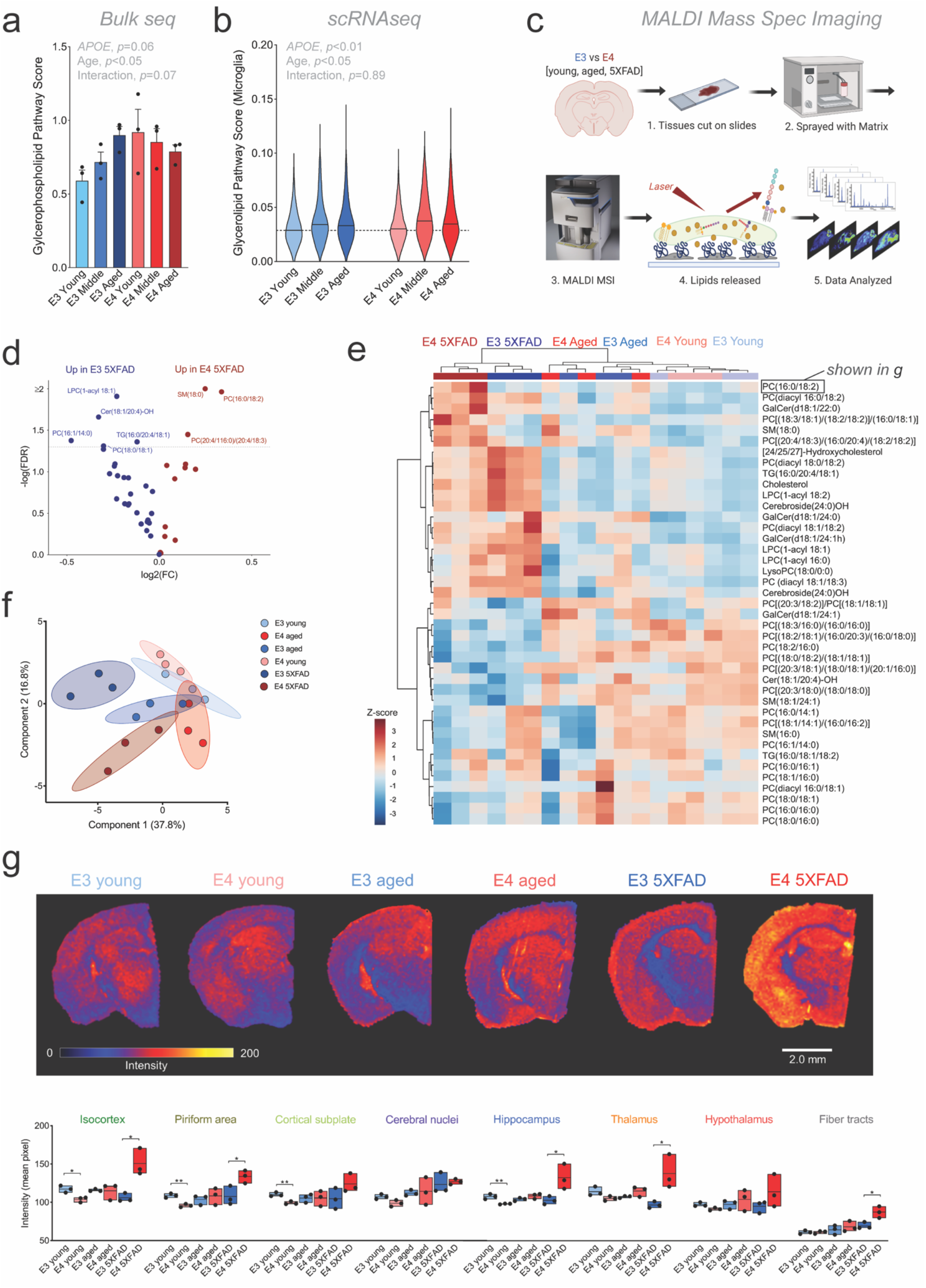
MALDI mass spec imaging reveals *APOE*- and region-specific changes in multiple lipid species. **a-b**, Expression of glycerophospholipid pathway genes increase with age in whole brain tissue (**a**, ‘*bulk*’) and is highest in aged E4 microglia (**b**, ‘*scRNAseq*’). **c**, Experimental workflow for matrix assisted laser desorption ionization (MALDI) mass spectrometry imaging (MSI). **d**, Volcano plot of targeted lipid species highlights changes in select phosphatidylcholine, sphingomyelin, ceramide and triacylglycerol. **e**, PCA plot of MALDI MSI detected lipids shows clear separation of E3 5XFAD and E4 5XFAD brains. **f**, Heatmap of quantified lipid species (average values across all regions) shows clear clustering by age and amyloid expression, with distinct separation of E3 5XFAD and E4 5XFAD brains. Brackets include multiple possible fatty acid chain lengths and/or double bond positions. **g**, Regional intensity of an example lipid from “e” (Phosphatidylcholine (16:0/18:2)). *Top*) Scans show spatial distribution of lipid across coronal brain sections. *Bottom*) Average pixel intensity across each brain region for PC(16:0/18:2). *n* =3 per group, \**p*<0.05, \*\**p*<0.01, multiple comparisons ANOVA. Regional data for all scanned lipids can be found in Table 2.

In order to validate the gene signatures implicating lipid dysregulation, we turned to MALDI MSI to generate qualitative, spatially resolved measures of targeted lipid species (**Fig. 8c**). Following fine-spatial scans of coronal sections, we assigned each MALDI MSI pixel to one of the anatomically assigned ST clusters. The overall lipid profiles showed clear heterogeneity, with samples generally clustering well by anatomical region (**Extended Data Fig. 8b**). The primary exception was the ‘disease associated’ area found almost exclusively in the E4 5XFAD brain. This region did not clearly cluster with itself nor with any other specific anatomical region, suggesting widespread dysregulation of lipid metabolism (**Extended Data Fig. 8b**).

The concentrations of multiple lipid species were altered in the E4 5XFAD brain relative to other groups, including multiple phosphatidylcholines (PCs) (**Fig. 8d-e**). Clustering analyses showed distinct separation of the E3 5XFAD and E4 5XFAD brains relative to other groups, and to each other (**Fig. 8f**). Interestingly, many of the observed age-, amyloid-, and *APOE*-associated changes in lipid concentrations were regionally specific (**Table 2**). For example, the PC most increased in the E4 5XFAD brain relative to E3 5XFAD (PC (16:0/18:2)) showed dramatic changes in the isocortex, hippocampus and thalamus, but no difference in the piriform area, cortical subplate and hypothalamus (**Fig. 8g**). Together, these results show that the transcriptional signatures implicating dysregulation of lipid metabolism in the E4 5XFAD brain are validated by alterations in multiple lipid species, in particular PCs.

## Discussion

It is increasingly appreciated that chronic neuroinflammation and metabolic dysfunction are early and prominent actors over the course of Alzheimer’s disease^42–44^. Notably, these two features are innately linked through the concept of immunometabolism^11,12^. Microglia are highly metabolically active cells^45^ that play a central role in maintaining CNS immune homeostasis, and the majority of genetic risk factors associated with late-onset AD are highly or specifically expressed in this cell type^46^. Many of these – including the strongest genetic risk factor for late onset AD, *APOE* – are thought to integrate metabolic inputs with downstream inflammatory signaling^47–49^. Here, we set out to systematically study the impact of *APOE* genotype across age, inflammatory challenge, and in response to amyloid using an integrative multi-omic approach. Collectively, our findings implicate *APOE4* as a driver of a dysfunctional immunometabolic response across each condition.

First, our ‘bulk’ tissue sequencing highlighted brain-wide changes in multiple immune and metabolic pathways similar to previous studies^28,30^. Though these data pointed toward E4-associated increases in metabolic, cytokine/chemokine, and complement pathways across the whole tissue landscape, the cell-specific changes potentially driving these bulk responses remained unknown. Therefore, we leveraged a scRNAseq approach using the same tissue samples analyzed for ‘bulk’ sequencing. In doing so, we identified a unique enrichment of a population of microglia, Mi_6, that predominates in the aged E4 brain. Differentially expressed biomarkers for this subcluster were enriched for genes involved in lipid metabolism and the innate immune response, as well as markers associated with DAM staging of microglia. Canonically, these populations emerge in response to neurodegenerative insults such as amyloid pathology, demyelination, or phagocytosis of apoptotic neurons^17,18^. It is therefore striking that a similar population of microglia (Mi_6) appears in aged E4 brains even while they lack discernible pathology.

KEGG pathway analyses conducted across microglia revealed terms of “Alzheimer’s disease”, “cholesterol metabolism”, and “HIF1α signaling” for the E4-enriched Mi_6 subset. Interestingly, *Hif1α* itself is a DAM gene, and many other important DAM genes (ex. *Spp1, Igf1*) are also HIF-responsive genes ^17,18,50^. Several recent reports have also demonstrated upregulated HIF1 signaling in AD, and collectively point to concurrent activation of both *Hif1α* and OxPhos gene expression as a common feature of amyloid-responding microglia in both humans and mice^37,51,52^. In line with this, our data also demonstrate that *Hif1α* was a predicted transcription factor enriched in Mi_6. *Hif1α* regulon activity correlated with a cell’s DAM score, a finding that may in part explain the high glycolysis in this subcluster. Additionally, in a reduced model system, E4 primary microglia had significantly higher gene expression of *Hif1α* compared to E3 microglia. Further, when we examined microglia harvested from E3 or E4 mice that received systemic LPS administration, the E4-LPS enriched Mi_8 and Mi_11 clusters both showed increased *HIF1α* activity and the highest glycolysis gene expression. Prior reports demonstrate that E4 is consistently tied to increased pro-inflammatory cytokine production after LPS stimulation in both humans^53^ and mice^54^. Bolstered by our current findings, we propose that this exaggerated inflammatory response, either due to chronic aging or acute pro-inflammatory exposure, may be a consequence of the unique E4-driven metabolic phenotype observed here across multiple paradigms.

Supporting this E4-associated bias toward *Hif1α* activation at the transcriptional level, we observed multiple functional indices of altered metabolism. For example, Seahorse analysis revealed increased aerobic glycolysis and decreased maximal mitochondrial respiration in E4 primary microglia compared to E3. Further, targeted metabolomics showed that E4 microglia display a marked accumulation of succinate and lactate, and a decrease in the anti-inflammatory metabolite itaconate. These findings partially align with prior work showing region-dependent alterations in mitochondrial respiration in the E4 mouse brain^30^ and reduced respiration and reduced glycolysis in human induced microglia-like cells (iMGLs) when edited from E3/E3 to E4/E4^55^. While E3 microglia responded to a pro-inflammatory stimulus by increasing mitochondrial ATP production, E4 microglia instead increased glycolytic ATP production. Increased mitochondrial respiration supports the switch to anti-inflammatory phenotypes whereas increased aerobic glycolysis enforces a pro-inflammatory phenotype^11,12^ Thus, the decreased maximal respiration and reliance on glycolysis for ATP production in response to pro-inflammatory challenge indicates that E4 microglia may be limited in their ability to engage mitochondrial ATP production, precluding effective tissue repair responses by preventing the switch to an anti-inflammatory phenotype. Collectively, our findings suggest that E4 microglia are predisposed to a pro-glycolytic, pro-inflammatory phenotype which they are then unable to resolve via metabolic reprogramming, setting up a situation conducive to chronic neuroinflammation.

We aimed to complement our bulk and scRNAseq data with a spatially resolved profile of the aging *APOE* transcriptome. ST highlighted the cortex and hippocampus as particularly vulnerable to E4-associated changes, which was exacerbated in the presence of amyloid. Specifically, *APOE4* appeared to exaggerate transcriptional ‘activation’ of microglia and was uniquely associated with alterations in lipid metabolism pathways in plaque-dense microenvironments. Interestingly, Fernández-Navarro *et al* identified gene expression changes in the hippocampus of 3xTg AD mice that pointed to altered metabolism^56^, including an upregulation of *Lpl* similar to our findings in both Mi_6 of aged E4 brains and the cortex of E4 5XFAD brains by ST. Using App^NL-G-F^ mice, Chen *et al* defined a plaque-induced gene signature (PIG) as well as an oligodendrocyte induced gene signature (OLIGs) comprised of genes responsible for remyelination^16^. Strikingly, we found that our magenta ‘microglial activation’ module strongly overlapped with the PIG signature, being lowest in E3 young, increasing with E4 and age, and highest in E4 5XFAD. This may suggest an exaggerated response to amyloid in E4 brains marked by increased microglial activation, in line with previous studies that have demonstrated increased pro-inflammatory responses in E4 microglia^54,57,58^. Since the PIGs signature is thought to represent dysregulated complement activation^16^, it is interesting to note that ApoE limits complement activation by forming a complex with C1q, but that the isoforms of ApoE have different binding profiles^59,60^. Thus, it is conceivable that the PIG^high^/OLIG^low^ profile seen in the E4 brain could lead to increased complement activation, aberrant pruning of synapses, and/or an imbalance in axon myelination, thereby feeding forward into a vicious cycle that propagates neuroinflammation and impaired lipid recycling.

Lipidomic analyses of the postmortem human AD brain have noted changes in brain lipids during the course of the disease^61^. Recent work extends these findings to include E4-associated decreases in several phospholipid species^62,63^ and isoform-specific microglial responses to ApoE-containing, phospholipid-rich lipoproteins^64^. While brain lipidomic profiling typically relies on tissue homogenates from preselected regions, we here leveraged MALDI MSI to simultaneously quantify multiple lipid classes across entire intact brain tissue sections. This allowed us to identify clear regional patterns of lipid expression that were substantially disrupted in the ‘disease associated’ cortical areas found primarily within the E4 5XFAD brain in our ST analyses. These ‘disease-associated’ brain regions had a gene signature that clearly mapped back to microglia (specifically Mi_6) in our scRNAseq dataset and was characterized by dramatic upregulation of DAM genes. Further, when we used the ST data to correlate our whole transcriptome profiles with amyloid plaque intensity on a spot-by-spot basis, we discovered a unique E4 signature that was highlighted by changes almost exclusively in pathways related to lipid metabolism.

Spatially-resolved quantification of lipids via MALDI MSI showed clear separation between the overall E3 and E4 brain lipidomes in aged mice, and more so in the amyloid overexpressing background of the 5XFAD brain. Specifically, they highlight *APOE*- and amyloid-associated decreases in a number of PCs, a class of phospholipid linked to memory decline^65^. While a handful of PCs were highest in the E4 5XFAD brain, the majority of PCs were present in lower concentrations relative to the other groups, consistent with findings from the human brain^62^. Regional segregation of lipid concentrations revealed that for most PCs, *APOE*-dependent changes in lipid concentrations were more striking in the isocortex, hippocampus, and fiber tracts relative to other brain regions. Strikingly, the top six PCs identified in our clustering analysis – where the highest concentrations were typically seen in the E4 5XFAD brain – are precursors of common phospholipid oxidation products, namely PCs having arachidonic acid (20:4) and linoleic acid (18:2). The multiple double bonds in these PCs are very labile to reactive oxygen species, with their oxidized forms being highly proinflammatory and associated with impaired mitochondrial ETC activity^66–69^. We also noted *APOE*- and amyloid-associated changes in cholesterol and several triglyceride (TG) species. These findings are intriguing as cholesterol esters and TGs are typically stored in intracellular lipid droplets (LDs), and recent studies have highlighted a role for glial LDs in aging and neurodegeneration, with E4 generally associated with LD accumulation and related metabolic disruptions^70–76^. Together, our bulk, scRNAseq, ST, and MALDI MSI data suggest that E4 predisposes AD-vulnerable brain regions to neurodegeneration through a metabolism-centered mechanism, perhaps owing to its altered binding profile for lipids and their receptors^77,78^.

Future work will help clarify the contribution of this *APOE*-associated perturbed immunometabolism in human AD, as aspects of the disease are not fully captured by mouse models. However, our use of humanized *APOE* TR mice^79–82^ not only allowed us to harvest fresh, high-quality tissue for scRNAseq and ST, but also appears to substantially bridge the gap between human and mouse studies, as we observe greater overlap with human transcriptomics studies compared to previous mouse studies that lacked human *APOE*. While some pathways altered by *APOE* were shared by multiple cell types (ex. KEGG “metabolic pathways”), many of E4’s effects on gene expression were unique to individual cell types and specific subpopulations. These single-cell differences appear important in understanding cross-species relevance. Serrano-Pozo *et al* recently re-analyzed publicly available data sets from large-scale bulk brain RNAseq of human AD patients to identify a transcriptional signature associated with E4 carriage^33^. Interestingly, when we compared this gene list to our scRNAseq data, over 50% of the pro-inflammatory and phagocytic genes upregulated in the brains of E4+ individuals with AD were significantly downregulated in young E4 microglia. This ‘flip’ in E4 microglia gene expression from lower in young to higher in aged was observed across many other gene lists from both mouse models of AD and human AD microglia studies, highlighting an age x *APOE* interaction in microglia programming. Finally, Patel *et al* recently analyzed fresh human surgical tissue from tumor or epileptogenic adjacent regions using scRNAseq. In line with our findings, they identified several WGCNA modules associated with age, sex and *APOE4* that were enriched for genes involved in lipid and carbohydrate metabolism^83^, a finding also reflected at the protein level by several large proteomics studies implicating immunometabolism in cortex^7,84,85^, CSF^7,86,87^, plasma^88^, and in isolated microglia^89^.

Collectively, our data suggest a potential scenario where metabolic dysfunction caused by APOE4 gives rise to chronic neuroinflammation, linking two phenomena consistently tied to AD with the strongest genetic predictor of the disease. These E4-associated immunometabolic disturbances appear intricately connected to aging and amyloid plaques, with the potential to exacerbate these pathological features and propagate synaptic loss through mechanisms of aberrant microglial activation and lipid dysregulation. This is especially true in the hippocampus and cortex, which were found to be uniquely vulnerable to E4’s immunometabolic reprogramming per our regional analyses. Many potentially modifiable AD risk factors such as obesity, diabetes, and physical inactivity also converge on immunometabolic pathways, as do other prominent genetic risk factors such as *TREM2*^90^, *CLU*^91,92^, and *BHLHE40*^93^. Thus, viewing Alzheimer’s disease through the lens of immunometabolism holds promise to fuse these seemingly disparate risk factors into a comprehensive mechanism whereby impaired microglial metabolism triggers chronic neuroinflammation, sparking the neurodegenerative cascade. Accordingly, therapies which target metabolism and inflammation in tandem may hold greater therapeutic promise in the treatment and prevention of AD.

## Methods

### Human *APOE* Mice

Human *APOE* ‘targeted replacement’ (TR) mice homozygous for *APOE3* or *APOE4* were employed across all experiments in the current study. In these “knock-in” mice, the coding region (exon 4) of mouse *Apoe* locus was targeted and replaced with the various human *APOE* alleles. Thus, human *APOE* expression remains under control of the endogenous mouse *Apoe* promoter, resulting in a physiologically relevant pattern and level of human ApoE expression^79–82^. Mice used for the bulk tissue RNAseq and scRNAseq portion of the study were female aged 3 months (young), 12 months (middle aged), or 24 months of age (aged). Mice used in the ST and MALDI MSI experiments were female mice aged 3 or 24 months of age, or female E3FAD or E4FAD mice (homozygous *APOE* TR mice crossed to the 5XFAD strain) 12 months of age. Mice used for the LPS study were females 12 months of age and intraperitoneally injected with saline or LPS (5mg/kg bodyweight) 24 hours prior to brain dissection. All mice were group housed in sterile micro-isolator cages (Lab Products, Maywood, NJ), and fed autoclaved food and acidified water ad libitum. Animal protocols were reviewed and approved by the University of Kentucky Institutional Animal Use and Care Committee.

### Cell culture

Primary mixed glial cultures were prepared from postnatal day 0–3 pups of mice homozygous for E3 or E4. The brain was surgically excised and meninges were removed from cortical tissue in ice cold dissection buffer (Hanks Balanced Salt Solution (Gibco # #14025-076) supplemented with 1% HEPES (Alfa Aesar #A14777), 1M sodium pyruvate (Gibco cat#11360-070), and 1% penicillin/streptomycin (Gibco # 15140-122). After dissection, isolated cortices were stored in a petri dish on ice containing growth medium (DMEM-F12 (Gibco #11320-033), 10% FBS (VWR# 97068-085), 1% penicillin/streptomycin). Tissue from 4-5 pups of the same genotype were pooled. Cortices were finely minced then transferred to a 15ml conical tube and dissociated with 5ml 0.25% trypsin-EDTA (Thermo #25200-056) for 25 minutes in a 37°C water bath with gentle agitation. An equal amount of growth medium was added to neutralize trypsin and the tubes were centrifuged at 300 × *g* for 5 min. After removing the supernatant, the tissue was washed three times with 2ml of warm HBSS. The tissue was then triturated in 10ml of warm growth medium and passed through a 70μm cell strainer (VWR #10199-657) to remove large particulates. Warm growth medium was added to a final volume of 10mL per mouse brain collected and seeded in T75 flasks (USA Scientific #658-175) incubated at 5% CO2 37°C. Medium was replaced with fresh growth media after 24hr. At 7 DIV medium was replaced with fresh growth medium supplemented with 10% L929 cell-conditioned medium (LCCM). LCCM contains growth factors including macrophage colony-stimulating factor 1 (M-CSF) that encourage microglial differentiation and proliferation. To prepare the LCCM, L929 cells (ATCC) were grown in DMEM/F12 with 10% FBS and 1% penicillin/streptomycin and the conditioned medium was harvested before passaging every 7 days, at which point it was centrifuged at 300 × *g* for 5 min and the supernatant sterile filtered through a 0.20μm vacuum filter and stored at −80C. Peak microglial confluence in the primary mixed glial cultures typically occurred around 12-14 DIV, at which point the flasks were shaken at 240rpm for 2 hours. Supernatant containing detached microglia was collected and centrifuged at 300 × *g* for 5 min. Cells were then resuspended and plated in supplemented growth medium incubated at 5% CO_2_ 37°C.

For cytokine stimulation experiments, cells were stimulated with a pro-inflammatory cocktail of 20ng/ml interferon-γ (IFNγ, R&D Systems #485-MI-100) and 50ng/ml tumor necrosis factor α (TNFα, R&D Systems #410-MT-025).

### Seahorse extracellular flux analysis

The Seahorse XF96 Glycolytic Rate Assay (Agilent) and Mitochondrial Stress Test (Agilent) were performed on E3 and E4 primary microglia to measure glycolysis and mitochondrial respiration, respectively. The Seahorse ATP Rate Assay (Agilent) was performed to measure the relative contributions of glycolysis and OxPhos to ATP production. Cells were seeded onto Seahorse XF96 tissue culture microplates (Agilent #101085-004) at a density of 3×10^4^ cells/well in supplemented growth medium (detailed above) and incubated at 5% CO_2_ 37°C. 12 hours prior to the start of the assay, cells were stimulated with pro-inflammatory (IFNγ + TNFα) cytokines as described above. The Seahorse Glycolytic Rate Assay (GRA), Mitochondrial Stress Test (MitoStress), and ATP Rate Assay were performed according to manufacturer’s instructions using DMEM-based medium containing 10mM glucose, 2mM glutamine, 1mM pyruvate, pH 7.4. For the GRA, plate was measured under basal conditions followed by serial addition of (A) rotenone and antimycin A (0.5μM) and (B) 2-deoxyglucose (50mM). For MitoStress, plate was measured under basal conditions followed by serial addition of (A) oligomycin (1μM), (B) FCCP (2.0μM), and (C) rotenone and antimycin A (0.5μM). For ATP Rate Assay, plate was measured under basal conditions followed by serial addition of (A) oligomycin (1.5μM) and (B) rotenone and antimycin A (0.5μM). Data were normalized to cell count using the automated Seahorse XF Imaging and Normalization System (Agilent) which utilizes 2μM Hoescht 33342 to label and count cell nuclei. Data were analyzed using Seahorse Wave v2.6 software (Agilent).

### Metabolomics

Primary microglia were plated at 7×10^6^ cells/well in 6-well plates (VWR #10062-894) and incubated at 5% CO_2_ 37°C. Upon reaching confluence, cells were removed from the incubator washed with warm 0.9% NaCl solution. Culture plates were placed on a bed of crushed dry ice and 1mL of ice cold 50% methanol (HPLC-grade, Sigma #A456-4) was added to quench cellular metabolic activity followed by a 10 minute incubation at −80°C to ensure cell lysis. After removing from the freezer, cells were detached with a cell scraper (VWR #10062-906) and the entire contents collected into a microcentrifuge tube, vortexed briefly, and placed on ice until all samples were collected. The tubes were then placed on a Disruptor Genie Cell Disruptor Homogenizer (Scientific Industries) for 5 min at 3,000 rpm. Tubes were then centrifuged at 40,000 × g for 10 min at 4 °C. The supernatant containing polar metabolites was isolated to a new tube and stored at −80C, and the resulting pellet was briefly dried at 10^−3^ mbar using a CentriVap vacuum concentrator (LabConco) to evaporate remaining methanol, followed by determination of protein content via BCA assay (ThermoFisher #23225) to normalize metabolite concentrations to total protein amount of each sample. The supernatant fraction containing polar metabolites was thawed gently on ice and dried at 10^−3^ mbar followed by derivatization. The dried polar metabolite pellet was derivatized by a two-step methoxyamine protocol first by addition of 70μL methoxyamine HCl (Sigma-Aldrich #226904-5G) in pyridine (20 mg/mL; Sigma-Aldrich #TS25730) to each pellet followed by 90 min dry heat incubation at 30°C. Samples were then centrifuged at 40,000 × g for 10 minutes after which 50μL of each sample was transferred to an amber V-shaped glass chromatography vial (Agilent #5184-3554) containing 80μL N-methyl-trimethylsilyl-trifluoroacetamide (MSTFA; ThermoFisher #TS48915) and gently vortexed followed by 30 min dry heat incubation at 37°C. The samples were allowed to cool to room temperature then analyzed via gas chromatography mass spectrometry (GCMS). Briefly, a GC temperature gradient of 130°C was held for 4 min, rising at 6°C/min to 243°C, rising at 60°C/min to 280°C and held for 2 min. Electron ionization energy was set to 70eV. Scan (m/z: 50–800) and full scan mode were used for metabolite analysis, and spectra were translated to relative abundance using the Automated Mass Spectral Deconvolution and Identification System (AMDIS) software with retention time and fragmentation pattern matched to FiehnLib library with a confidence score of > 80. Chromatograms were quantified using Data Extraction for Stable Isotope-labelled metabolites (DExSI) with a primary ion and two or more matching qualifying ions. Metabolomics data was analyzed using the web-based data processing tool Metaboanalyst ^94^.

For stable-isotope resolved metabolomics, cells were washed with warm, sterile phosphate-buffered saline (PBS; Thomas #QZY-11666789001-4L) to remove traces of non-^13^C media and then incubated in glucose- and sodium pyruvate-free DMEM (Thermo #11966-025) containing 2mM GlutaMAX (Thermo #35050-061), 1% penicillin/streptomycin, and 10mM universally labelled ^13^C-glucose (Cambridge Isotope Laboratories # CLM-1396-PK) for two hours with pro-inflammatory stimulus (IFNγ + TNFα, as described above). After the two hours incubation metabolites were extracted from the cells and processed for GCMS as described above. Fractional enrichment was calculated as the relative abundance of each isotopologue relative to the sum of all other isotopologues.

### Quantitative PCR

E3 and E4 primary microglia were plated at 5×10^6^ cells/well in 6-well plates and RNA was extracted from the cells using the RNEasy Plus Mini Kit (Qiagen #74136) and converted to cDNA using High-Capacity RNA-to-cDNA kit (Thermo #4387406) according to manufacturer’s instructions. TaqMan chemistry was used for quantitative PCR with TaqMan probe targeting *Hif1a* (Thermo #4453320) and TaqMan Fast Advanced Master Mix (Thermo #4444556), PCR was performed on the QuantStudio 3 (Applied Biosystems) with default cycling parameters for this master mix (initial holds at 50°C for 2 minutes (UNG incubation) and 95°C for 20 seconds (polymerase activation) then 40 cycles of denaturation at 95°C for 1 second followed by annealing/extension at 60°C for 20 seconds). Data were analyzed using the ddCT method with 18s ribosomal rRNA (TaqMan assay id# Hs99999901_s1) as the reference gene.

### Brain single-cell suspension, cDNA library, and sequencing

Pooled brain tissue (n=3 per experimental group) was processed for ‘glia-enriched’ single cell suspensions as previously described^95^. Briefly, mice were anesthetized via 5.0% isoflurane before exsanguination and transcardial perfusion with ice-cold Dulbecco’s phosphate buffered saline (DPBS; Gibco # 14040133). Following perfusion, brains were quickly removed, pooled, and whole left hemispheres sans brainstem and cerebellum were quickly minced using forceps on top of an ice-chilled petri dish. Minced tissue from the 3 pooled hemispheres per group were immediately transferred into gentleMACS C-tube (Miltenyi #130–093-237) containing Adult Brain Dissociation Kit (ADBK) enzymatic digest reagents (Miltenyi #130–107-677) prepared according to manufacturer’s protocol. Tissues were dissociated using the “37C_ABDK” protocol on the gentleMACS Octo Dissociator instrument (Miltenyi #130–095-937) with heaters attached. After tissue digestion, cell suspensions were filtered through 70 μm mesh cell filters to remove debris following the manufacturer’s suggested ABDK protocol. The resultant suspension was sequentially filtered (x2) using fresh 30 μm mesh filters. Cell viability was checked using AO/PI viability kit (Logos Biosystems # LGBD10012). All cell suspensions were determined to have > 90% viable cells. Following viability and counting, cells were diluted to achieve a concentration of ∼ 1700 cells/μL in a 10μL total reaction volume. The diluted cell suspensions were loaded onto the 10X Chromium Connect automated cell portioning system. Sample libraries were constructed using Next GEM automated 3’ reagents (10X Genomics, v3.1) following manufacturer’s suggested protocol (#CG000286 Rev B). Final library quantification and quality check was performed using BioAnalyzer (Agilent), and sequencing performed on a NovaSeq 6000 S4 flow cell, 150 bp Paired-End sequencing (Novogene).

### scRNAseq data processing

After libraries were sequenced and quality control was performed, samples were aligned to the mm10 mouse reference genome using the Cell Ranger 6.0.2 pipeline. Each sample was aggregated using the cellranger aggr function to produce a raw UMI count matrix containing the number of reads for genes in each cell per sample. The expression matrix was loaded into R for further analysis and visualization using Seurat (v.4.1.0). Cells were then filtered to reduce the potential of including doublet and low-quality cells using the following criteria: 200 < nGene < 5000; 500 < nCount < 90,000; and percent.mito <30%. Feature counts were normalized using LogNormalize method with a scale factor of 10,000 (default option); and the effects of percent.mito were regressed out using the ScaleData method. A shared nearest neighbor (SNN) graph was constructed using FindNeighbors function with default parameters. Using the Louvain algorithm implemented in FindClusters function and the first 15 principal components (PCs), we identified 34 unique clusters.

To assign glial cell type identity to each cluster, we manually examined the expression levels of cell type-specific markers across each cluster using Partek software to identify clusters containing unique populations of different cell types. Canonical CNS cell type markers were compiled from^96–99^ and included: *Aldoc, Aqp4, Gja1, Aldh1l1, Gfap, Slc7a10, Sox9* (Astrocytes), *P2ry12, Tmem119, Aif1, Slc2a5, Trem2, Cx3cr1, Itgam, Gpr34, C3ar1, Csf1r, Fcrls* (Microglia), *Mgl2, Mrc1, Pf4* (Macrophages), *Mog, Opalin, Mag, Ermn, Cldn11* (Oligodendrocytes), *Pdgfra, Opcml, Tnr, Myt1* (Oligodendrocyte precursors), *Kl, Car12, Ttr* (Choroid plexus), *Ccdc153, Dnah11* (Ependymal), *Cd3d* (Lymphocytes), *Flt1, Emcn, Cldn5, Cdh5, Vwf, Tek, Cd34* (Endothelial), *Slc47a1, Mgp* (Vascular leptomeningeal), *Acta2, Bgn* (Vascular smooth muscle), *Vtn, Kcnj8* (Pericytes), and *Dcx* (Neuroprogenitors). This process resulted in stringent filtering of cells with ambiguous assignments (>1 cell-specific gene marker; likely ‘doublet’ and ‘triplet’ that slipped through the 10X ‘single cell’ droplet workflow), leaving a total of 39,475 cells within 24 carefully assigned glial clusters.

### Re-clustering of specific glial cell populations (ex. microglia)

We used the FindAllMarkers function to identify genes that act as markers for each cluster, using the Wilcoxon rank-sum test. A gene was considered the marker of a cluster if it had a Bonferroni-adjusted p-value <0.01 and an average log fold change >0.1. The data were further filtered to contain only astrocytes, microglia or other glial cell types using the markers described above. After re-clustering with a resolution of 0.1 and the first 15 PCs, we identified 11 microglia sub-clusters and 12 astrocyte sub-clusters. To perform differential expression analysis in the cell-specific datasets, we used Seurat’s FindMarkers function and performed Wilcoxon rank-sum tests. A gene was considered differentially expressed if it had a Bonferroni-corrected p-value <0.05 and a natural log fold change (logFC) >0.25.

### Pathway enrichment analyses of GO Terms

The Seurat function FindMarkers conducted the DEG analysis via grouping for comparison by clusters, *APOE* genotype, or age (min.pct was set as 0.25 and logFC.threshold was set as 0.25). The DEGs were selected if the adjusted p-value was less than 0.05 and the absolute value of log-fold change was higher than 0.1. Based on the identified DEGs, the enrichment analyses of GO terms (Biological Process (BP) were performed via Enrichr^100^ or the rWikiPathways R software package^101^ with cutoff by FDR-adjusted adjusted p values 0.05. The bar-plot functions from the software package with a color-blind-friendly color scheme were applied for the visualizations.

### Gene set enrichment analysis for metabolic pathways gene signatures

AUCell R software package (v.1.14.0) was applied for the identification of gene signatures at the single-cell level^34^. AUCell uses the “Area Under the Curve” (AUC) to calculate whether a critical subset of the input gene set is enriched within the expressed genes for each cell. AUC scores were calculated for each individual cell and distribution across cell populations of interest allowed for exploration of the relative expression of the gene signature. AUCell scores of seventy KEGG pathway gene sets associated with pathways of mammalian metabolism (https://www.genome.jp/kegg/pathway.html) were manually curated and applied to multiple datasets The average AUCell scores of each pathway were plotted as heatmaps using pheatmap R software (v.1.0.12) sorted by either cell subtypes and/or experimental groups.

### Gene-gene network analysis using WGCNA

Weighted gene co-expression network analysis (WGCNA) (v1.70-3)^102^ was used to identify gene modules and build unsigned co-expression networks, including both negative and positive correlations. Briefly, WGCNA constructs a gene co-expression matrix, uses hierarchical clustering in combination with the Pearson correlation coefficient to cluster genes into groups of closely co-expressed genes termed modules, and then uses singular value decomposition (SVD) values as module eigengenes (MEs) to determine the similarity between gene modules or calculate association each module with sample traits (ex. *APOE* genotype and treatment). The top 3,000 variable genes were selected to identify gene modules and network construction.

Soft power of 6 was chosen by the WGCNA function pickSoftThreshold. Next the function TOMsimilarityFromExpr was used to calculate the TOM (Topological Overlap Matrix) similarity matrix via setting power = 6, networkType = “signed”. The distance matrix was generated by subtracting the values from the similarity adjacency matrix by one. The function flashClust (v.1.01) was used to cluster genes based on the distance matrix, and the function cutreeDynamic was utilized to identify gene modules by setting deepSplit =3. Cytoscape (v.3.8.2) was applied for the gene-gene network visualization.

### Gene Score Plots

Pairwise differential expression analyses were then performed between E4 vs E3, aged vs young, aged vs middle, and middle vs young. For each gene within each differential expression analysis, a gene score was calculated to represent a combination of effect size and statistical significance of the differential expression. The gene score was calculated as the product of the log2 fold change (FC) and negative of the log-transformed false discovery rate (FDR), log2(FC)*-log10(FDR).

### Gene transcriptional regulatory network analyses using pySCENIC

For regulon identification, gene regulatory network analysis was performed using the pySCENIC software packages (v.0.11.2)^103^. The arboreto package is used for this step using the algorithm of GRNBoost2 (version 0.11.2) to identify the potential transcriptional factor (TF)-targets based on their co-expression with RcisTarget (version 1.12.0) for cis-regulatory motif enrichment analysis in the promoter of target genes (mm9-500bp-upstream-10species.mc9nr and mm9-tss-centered-10kb-10species.mc9nr databases provided in the pySCENIC package), and to identify the regulon, which consists of a TF and its co-expressed target genes. Correlations between a list of 1,390 human transcription factors (TFs) curated by Lambert *et al*^104^ and the genes in the expression matrix were evaluated, and co-expression modules with a minimum size of 20 genes were defined. Finally, for each regulon, pySCENIC uses the AUCell algorithm to score the regulon activity in each cell. The input for SCENIC was the n (genes) by n (cells) matrix obtained after filtering, and gene expression is reported in count units. Parameters used for running were specified as default options in the original pySCENIC pipeline. The cellular activity pattern of a predicted regulon can be binarized as being in an ‘on’ or ‘off’ state based on the bimodal distribution of a regulon’s AUCell values and visualized as a heatmap for identification of regulon clustering.

### Brain preparation for spatial transcriptomics

The mirroring hemisphere (right) from brains processed for scRNAseq (see Methods section “Brain single-cell suspension, cDNA library, and sequencing”) were immediately placed in OCT compound (Fisher HealthCare Tissue Plus O.C.T. Compound Clear 4585) and gently lowered into isopentane (Sigma-Aldrich 2-Methylbutane M32631) in a beaker surrounded by dry ice (isopentane chilled to approximately −70°C). Brains were submerged for 60 seconds, placed on dry ice, wrapped in aluminum foil, and stored at −80°C until sectioning. Prepared brain hemispheres were cryosectioned to 10 μm thick coronal sections at approximately Bregma −2.00 mm. Serial 10 μm sections immediately rostral and caudal to the section mounted on the Visium Spatial Gene Expression slide (10X Genomics) were collected for immunohistochemistry. Optimal tissue permeabilization time was determined using the manufacturer’s optimization protocols (10X Genomics, Visium Spatial Tissue Optimization), and accordingly, experimental tissues were permeabilized for 18 min for Visium Spatial Gene Expression analysis. Prior to library preparation, tissue sections were methanol-fixed, stained with hematoxylin and eosin (H&E), and imaged on a Nikon NiU microscope with Fi3 color camera. Sections were then permeabilized and processed to obtain cDNA libraries, which were subsequently prepared according to the manufacturer’s protocol (https://support.10xgenomics.com/spatial-gene-expression/library-prep). Final library quantification and quality check was performed using BioAnalyzer (Agilent), and sequencing performed on a NovaSeq 6000 S4 flow cell, 150 bp Paired-End sequencing (Novogene).

### Spatial transcriptomics data processing

Raw FASTQ data and H&E images were processed by the Space Ranger v1.3.0 (10X Genomics) pipeline. Illumina base call (BCL) files from the sequencing instrument were converted to FASTQ format for each sample using the mkfastq. Visium spatial expression libraries were analyzed with the count command. Image alignment to predefined spots was performed by the fiducial alignment grid of the tissue image to determine the orientation and position of the input image. Sequencing reads were aligned to the mm10 reference genome using STAR (v2.5.1b) aligner. Gene expression profiling in each spot was performed with UMI and 10X barcode information. The spots with gene expression data were analyzed with the Seurat package (v.4.1.0). Gene counts were normalized using ‘LogNormalize’ methods in Seurat. The top highly variable genes (n= 3,000) were then identified using the ‘vst’ method in Seurat. The number of RNA counts for each spot and the frequency of mitochondrial gene counts were regressed out in the scaling process. Six spatial transcriptomic datasets were merged and rescaled (E3 young, E4 young, E3 aged, E4 aged, E3 5XFAD, E4 5XFAD). Principal component analysis was performed using the top highly variable genes. For visualization, dimension reduction was performed using UMAP on the top 20 principal components were applied. Graph-based clustering based on the Louvain community detection algorithm was performed. Markers for each cluster were identified by Wilcoxon rank-sum tests for a given cluster vs. other clusters implemented in Seurat as a ‘FindAllMarkers’ function.

### Integrative analysis of amyloid plaque intensity and spatial transcriptomic data

The anatomical location of each cluster was visually identified by comparison with the Allen Mouse Brain Reference Atlas (https://mouse.brain-map.org/static/atlas). The region annotation information (ex. isocortex, fiber tracts, etc.) was integrated as spot metadata. Separately, the amyloid plaque (X-34 stained) images were prepared from a 10 μM section immediately adjacent (caudal) to the 10 μM section used for the ST data generation. The plaque image was resized to exactly match the same-section H&E image for the ST coordinates. Cropping and rotation were performed to overlap both images, and the color channels specifically addressing the plaque intensities (X-34, blue) were extracted using the Photoshop image analysis tool. The quantitative extraction of plaque intensity scores was performed using the Squidpy software package (version 1.0.0)^105^. The resulting plaque intensity score values were added as spot metadata for downstream analyses.

### Ligand–receptor cell-cell interactions

Cell-to-cell communication was identified by evaluating the expression of pairs of ligands and receptors within cell populations using the CellChat R software package (version 1.1.3)^106^. CellChat infers the biologically significant cell-cell communication by assigning each interaction with a probability value and performing a permutation test. CellChat models the probability of cell-cell communication by integrating gene expression with prior known knowledge of the interactions between signaling ligands, receptors, and their cofactors using the law of mass action. We examined the interaction among different cell types or microglia and astrocyte subtypes. The databases, including ‘Secreted Signaling’ provided by Cellchat, were used. To identify the ligand-receptor interactions of ApoE, a gene list of *APOE* and target receptors (ex. *Ldlr*) reported by Sheikh et al.^107^ was added to the receptor-ligand interaction database of CellChat. The ApoE-ApoE receptor list is included in Table 1.

### Matrix Assisted Laser Desorption Ionization (MALDI) mass spectrometry imaging (MSI)

Brain sections (10 μm) were mounted on glass slides and prepared for MALDI MSI (see Method section “Brain preparation for spatial transcriptomics”). Slides were prepared as previously described^108^. Briefly, after desiccation for one hour, slides were sprayed with 14 passes of 7mg/mL N-(1-Naphthyl) ethylenediamine dihydrochloride (NEDC) matrix (Sigma) in 70% methanol (HPLC-grade, Sigma) was applied at 0.06mL/min with a 3mm offset and a velocity of 1200mm/min at 30°C and 10psi using the M5 Sprayer with a heated tray of 50°C. Slides were used immediately or stored in a desiccator until analysis. For the detection of lipids, a Waters SynaptG2-Xs high-definition mass spectrometer equipped with traveling wave ion mobility was used as previously described^108^. The laser was operating at 2000 Hz with an energy of 300 AU and spot size of 50 μm at X and Y coordinates of 100μm with mass range set at 50 – 1000 *m/z* in negative mode. MALDI-MSI data files were processed to adjust for mass drift during the MALDI scan and to enhance image quality and improve signal-to-noise ratio using an algorithm available within the High-Definition Imaging (HDI) software (Waters Corp). To adjust for mass drift during the MALDI scan, raw files were processed using a carefully curated list of 20 MALDI NEDC matrix peaks (*m/z*) 26 small molecule MALDI peaks(*m/z*), and 24 lipid peaks(*m/z*). Files were processed at a sample duration of 10 sec at a frequency rate of 0.5 min, and an *m/z* window of 0.1 Da, using an internal lock mass of previously defined metabolite of taurine 124.007 *m/z* with a tolerance of 1amu and a minimum signal intensity of 100,000 counts. Data acquisition spectrums were uploaded to the HDI software for the generation of lipid images. Regions of interest (ROIs) were user defined by a blinded investigator using anatomical reference points based on the mouse Allen Brain atlas. For all pixels defined within a ROI, peak intensities were averaged and normalized by total ion current (TIC) and number of pixels.

## Acknowledgements

We thank Dr. Doug Harrison and Jim Begley at the University of Kentucky Arts & Sciences Imaging Center for their invaluable assistance with scRNAseq and ST analyses. We also thank Dr. Matthew Gentry for his continued support and guidance. Finally, we thank Dr. Tomoko Sengoku and Michael Alstott at the University of Kentucky Redox Metabolism Shared Resource Facility, supported by NCI Cancer Center Support Grant (P30 CA177558), for their assistance with the Seahorse assays.

## Author Contributions

LAJ and JMM designed the experiments. SL, NAD, JMM and LAJ analyzed the data and wrote the paper. NAD completed the metabolic analyses of microglia, including metabolomics, Seahorse assays, and RT-PCR. EJA and JLS performed tissue preparation, sectioning and staining for spatial transcriptomics and immunohistochemistry, respectively. CTS, CMF, AEW, GMS, SMM and HCW assisted with bulk and scRNAseq analyses. JMM and LAJ supervised scRNAseq and ST workflows, with technical assistance from JLS, AAG, and DSG. RCS oversaw MALDI mass spec imaging, with technical and analytical assistance from LRG, HAC and TRH. All authors read the paper and provided edits.

## Supplemental Figures

**Extended Data Fig. 1 (related to Fig. 1).**
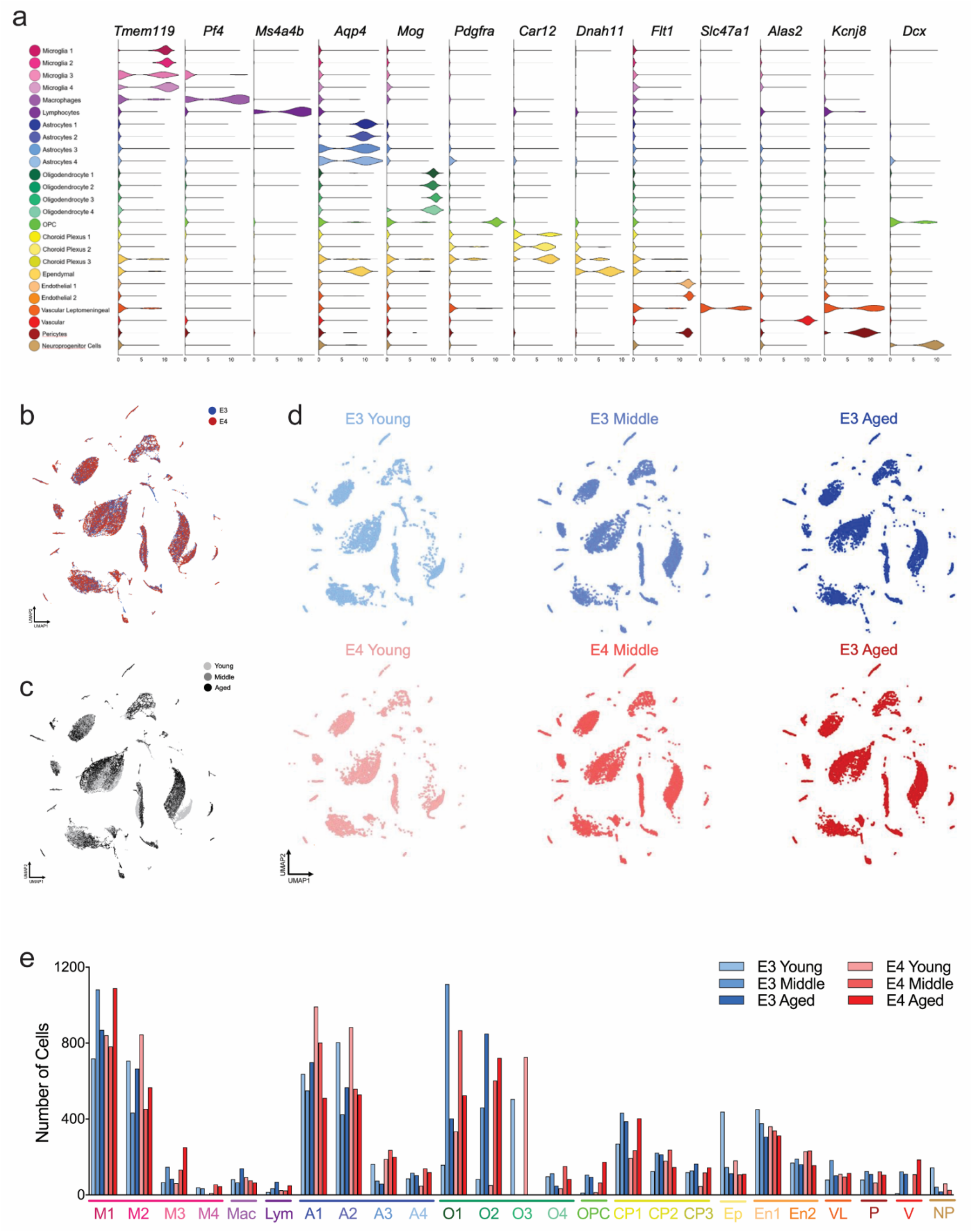
Cell-specific markers, UMAPs and cell numbers per cluster. a) Gene expression of cell-specific markers for each cluster. b-d) UMAP plots of all cells color coded by *APOE* genotype (b) or age (c), or split by experimental group (d). e) Total number of cells in each cluster.

**Extended Data Fig. 2 (related to Fig. 2).**
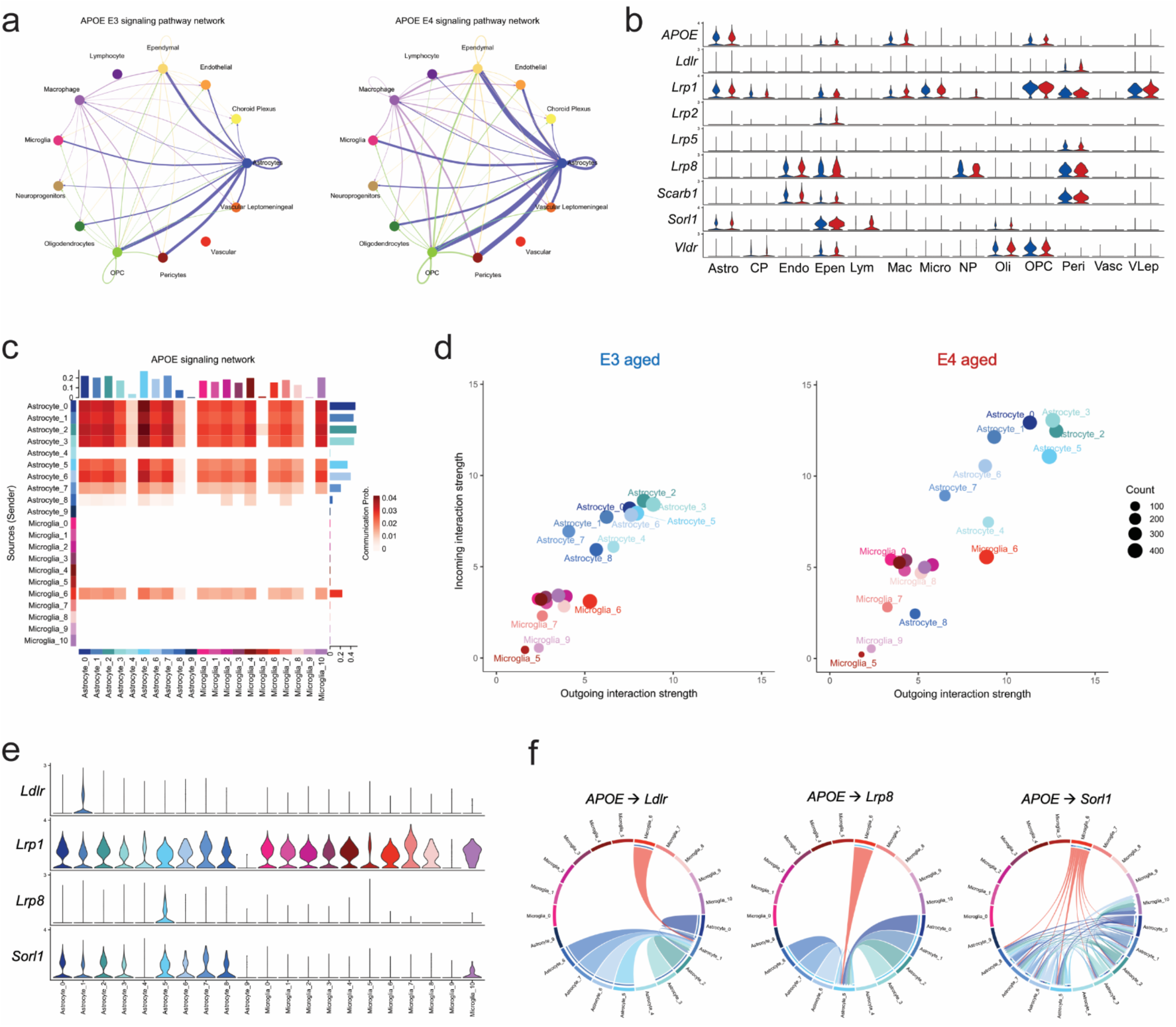
ApoE-ApoE receptor signaling pathway network analyses. a) Circle plot of signaling pathway networks for E3 (left) and E4 (right) cel types. Lines connecting cell types denote the aggregated cell-cell communication probabilities evaluated from the expression of the receptors, ligands, and cofactors in different types in the scRNA-Seq dataset. The width of lines corresponds to the signaling pathways’ relative interaction strength (aggregated communication probabilities). b) Expression of *APOE* and various ApoE receptors across all glial cell types in E3 (blue) and E4 (red). c) Heatmap of the ApoE-ApoE receptor signaling pathway network across the various astrocyte and microglia subclusters in the E3 and E4 cell datasets. Heatmap color represents communication probability, i.e. the probability of cell-cell communication by integrating gene expression with prior known knowledge of the interactions between signaling ligands, receptors, and their cofactors. Bars represent the sum of the communication probabilities of each subtype. X- and Y-axis indicate the targets (cells expressing receptor) and sources (cells expressing ligand) of the signaling communication. d) Interaction plots showing the outgoing (*APOE* expression) and incoming (ApoE receptor (Ldlr, Lrp1, etc.) expression) signal strength for each astrocyte and microglia subcluster in two sub-datasets (E3 and E4). Circle size represents the number of signaling pathways of *APOE* and its receptors among a specific cell subtype and the other cell subtypes. e) Subcluster gene expression of the four ApoE receptors expressed at detectable levels by astrocytes or microglia. f) Circos plots showing significant ApoE-receptor interactions within astrocyte and microglia subclusters for *Ldlr* (left), *Lrp8* (middle), and *Sorl1* (right). The outer circle color denotes the sending subcluster (*APOE* expressing), while the inner circle color denotes the receiving cluster(s) (a specific ApoE receptor expressing).

**Extended Data Fig. 3 (related to Fig. 3).**
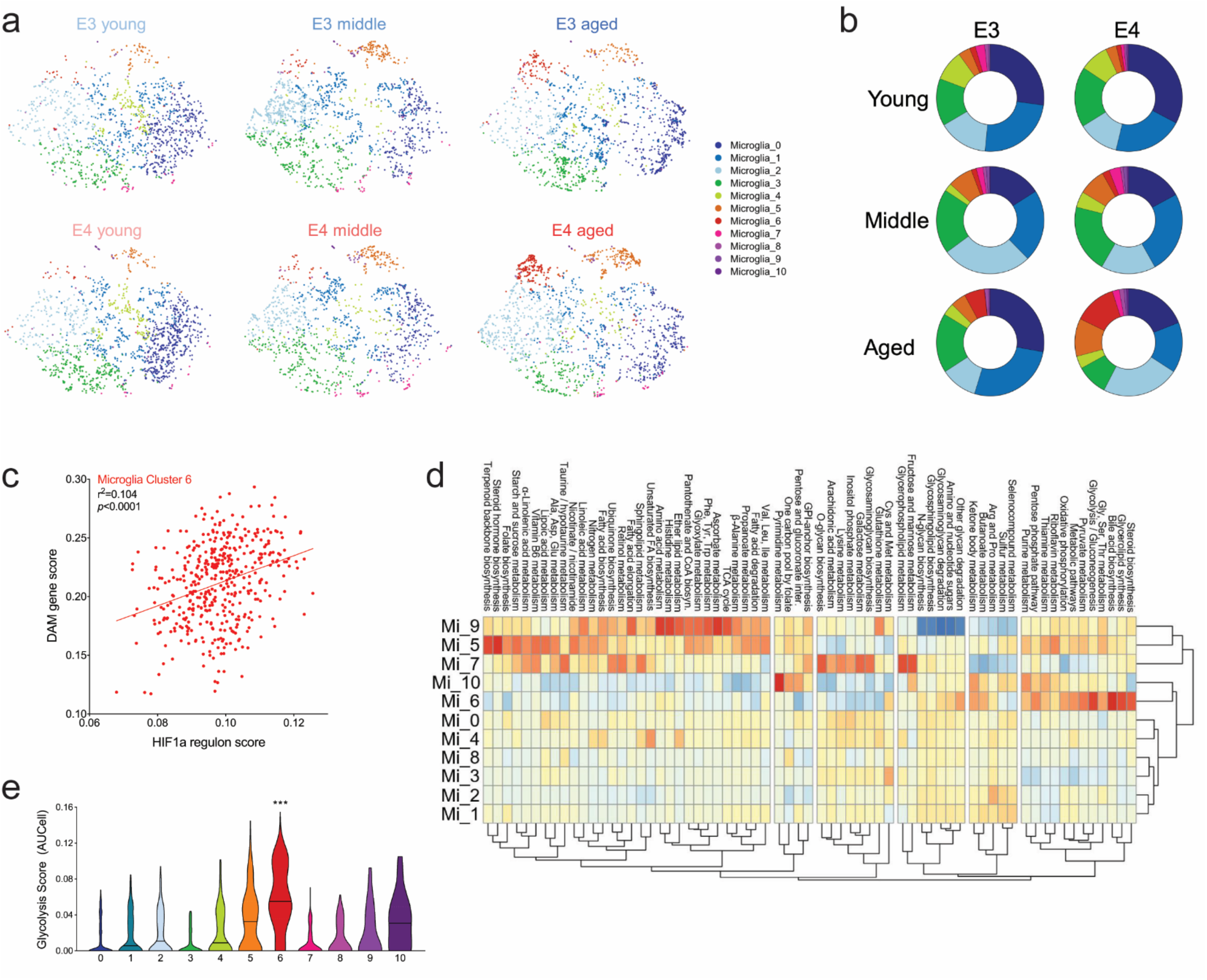
Cell distribution and metabolic gene expression in microglia subpopulations. a-b) Distribution of microglia across clusters. a) tSNE of microglia clusters separated by experimental group. b) Donut charts showing the distribution of E3 (left) and E4 (right) microglia within each cluster across ages. c) HIF1a regulon score correlates with DAM/MgND gene scores for microglia in cluster 6. d) Heatmap showing expression of KEGG metabolic pathways in each microglia cluster. e) Glycolysis pathway scores for each microglia cluster. ***, *p*<0.001 Mi_6 vs all other clusters.

**Extended Data Fig. 4 (related to Fig. 5).**
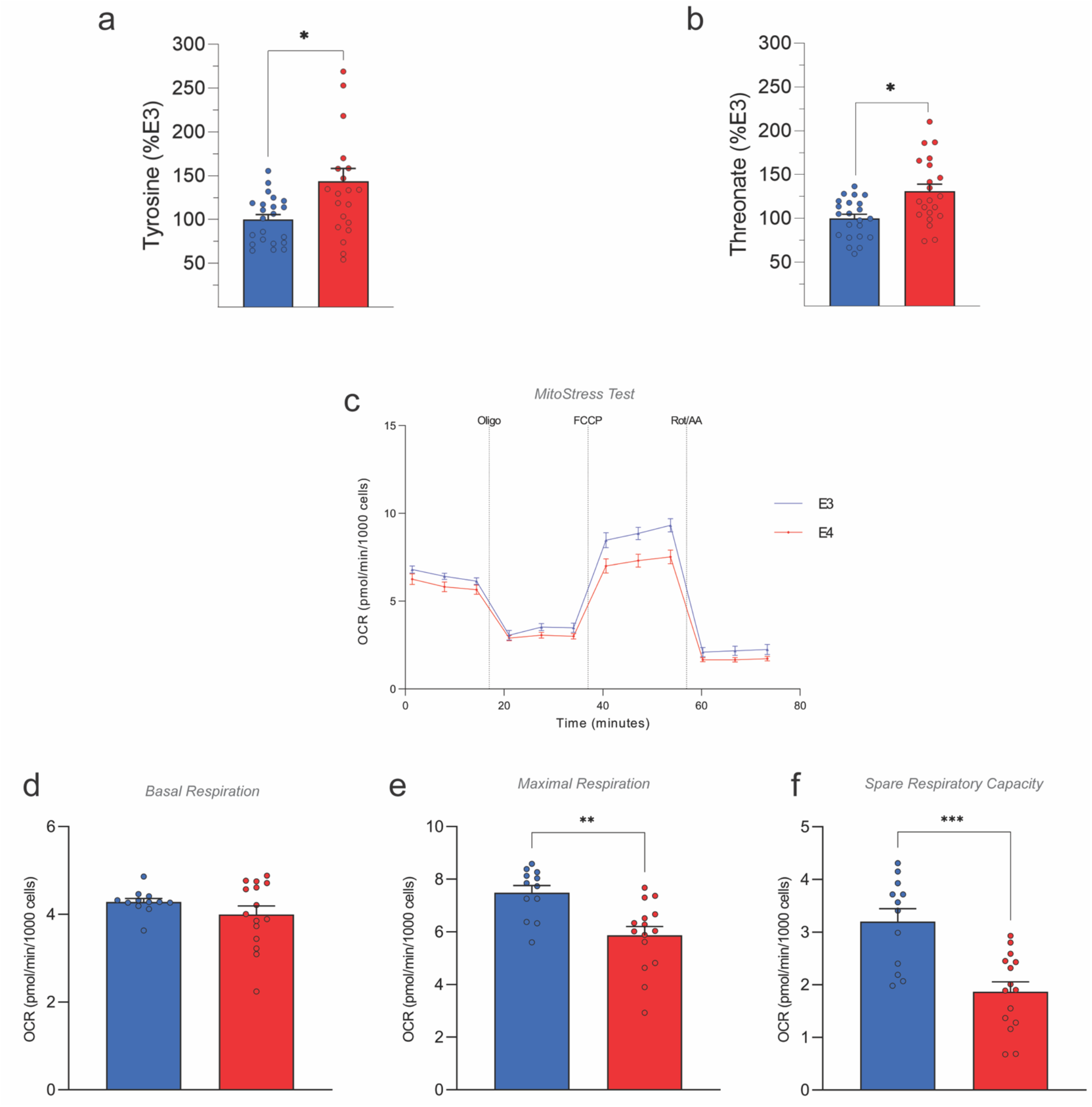
Targeted metabolomics and oxygen consumption rate in E3 and E4 microglia. a-b) Tyrosine and threonate concentrations from targeted metabolomics of E3 and E4 microglia. c-f) E3 and E4 microglia were assayed using the Seahorse Mito Stress Test (n=12-15 per group). c) Oxygen consumption rate (OCR) measured over time in E3 and E4 microglia. E3 and E4 microglia showed similar basal respiration (d) whereas E4 microglia showed the lowest maximal respiration (e) and spare respiratory capacity (f). *p<0.05, **p<0.01, ***p<0.001, two-tailed T-test.

**Extended Data Fig. 5 (related to Fig. 6).**
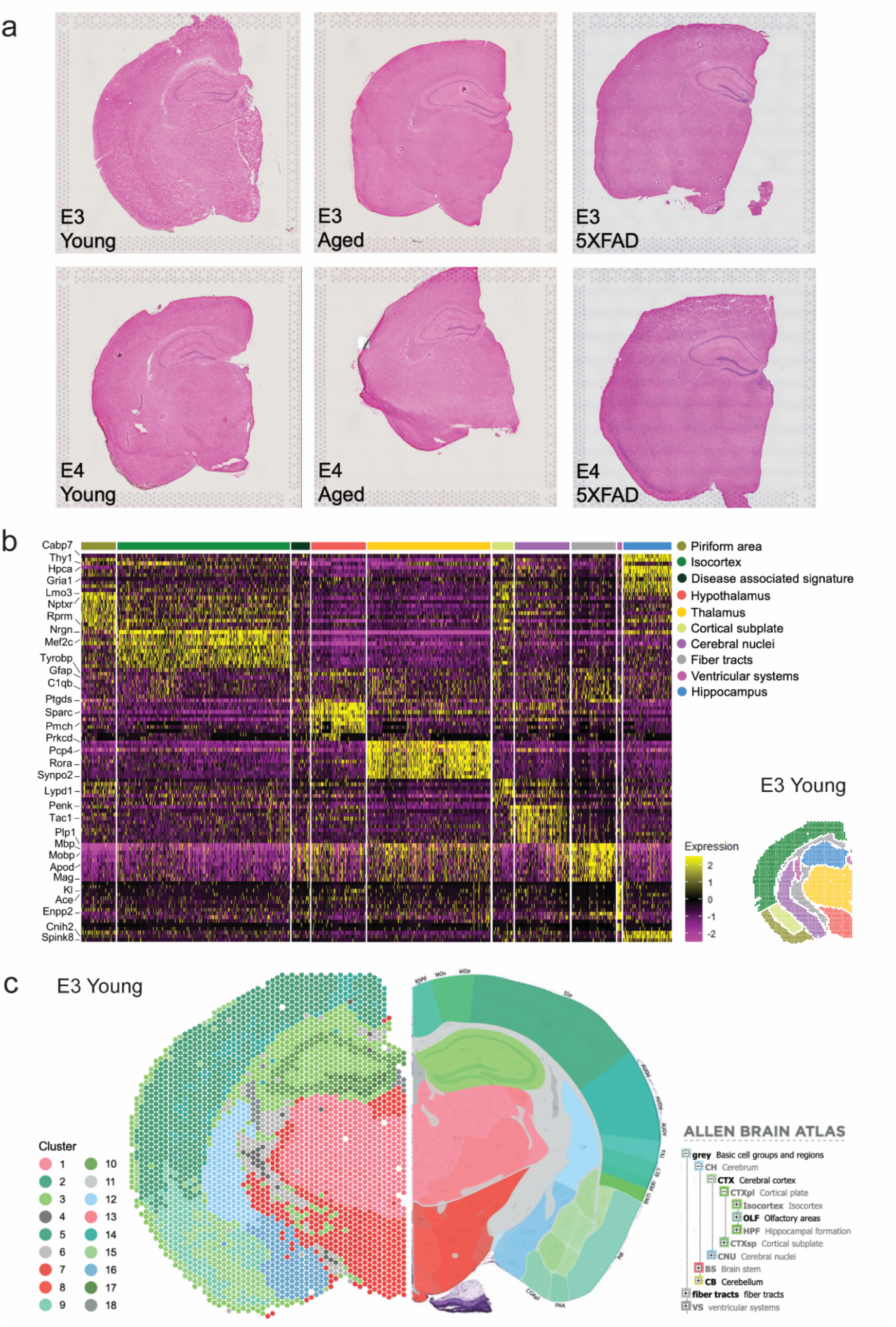
Spatial transcriptomics shows clear clustering based on brain anatomy, plus a distinct “Disease associated signature”. a) Hematoxylin and eosin (H&E) stain of coronal brain sections subjected to spatial transcriptomics (ST) analysis (Visium Spatial Gene Expression, 10X Genomics). b) Heat map showing expression of genes (rows) used to define the clustering of each ST spot (columns) within 9 anatomically-defined brain regions, plus the distinct “Disease associated signature” cluster. Various representative ‘biomarker’ genes for each cluster are shown on the left. Colored bars denote the anatomical region assigned to each cluster. Inset shows a representative brain (E3 young) color coded to reflect the assigned anatomical regions (white spots denote unassigned ‘border regions’). c) Recoloring of the same 18 unbiased clusters from Figure 6 (left) to show close correspondence with anatomical brain regions of the Allen Brain Atlas (right). Representative brain shown is the ‘E3 young’.

**Extended Data Fig. 6 (related to Fig. 6).**
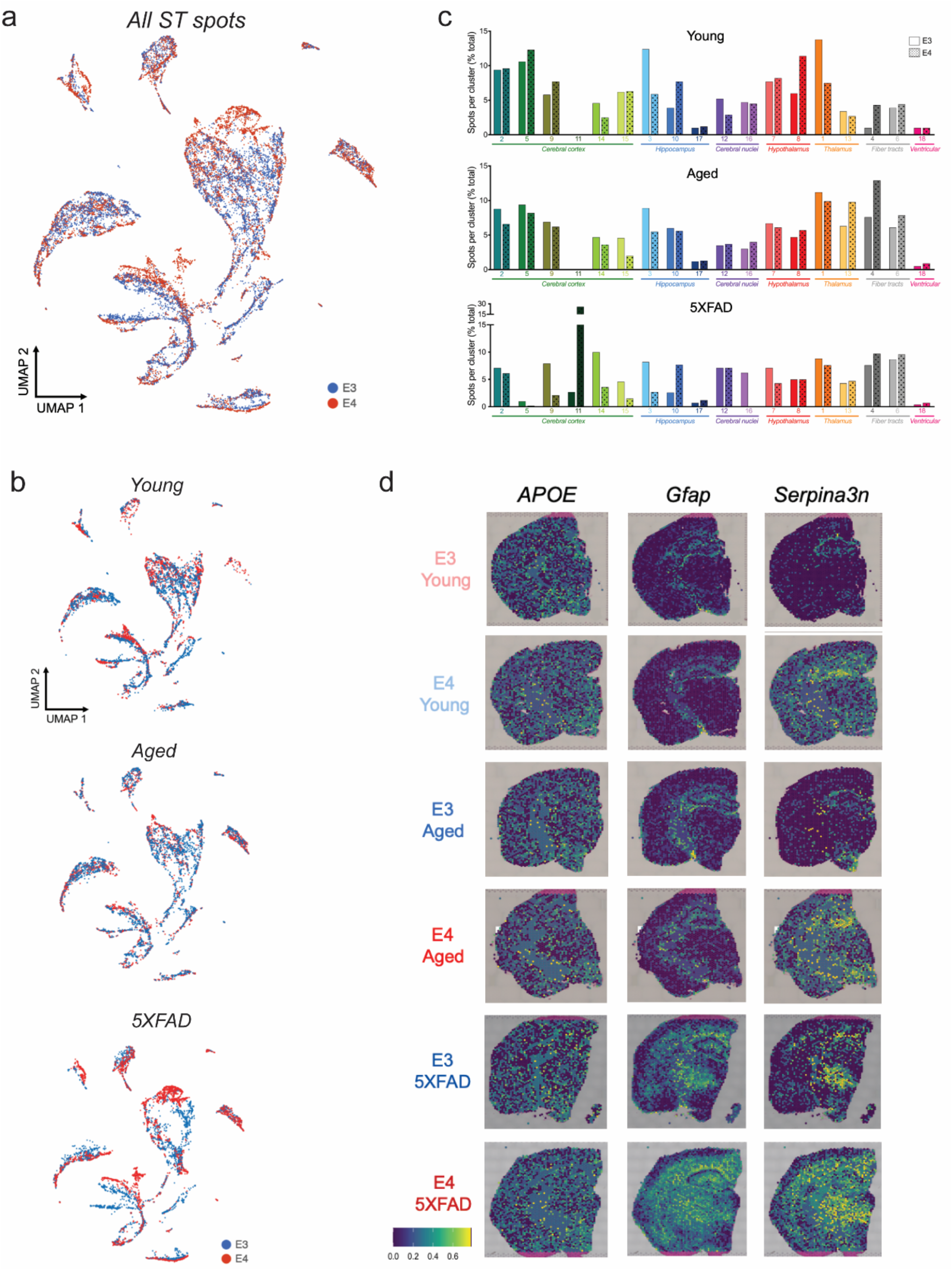
Spatial transcriptomics UMAPs and spot number per cluster. a) UMAP showing all 79,980 spatial transcriptomic (ST) spots, colored by *APOE* genotype (E3 blue, E4 red). b) UMAPs showing ST spots from young, aged, or 5XFAD mice with E3 (blue) or E4 (red). c) Number of spots within each cluster in young, aged, or 5XFAD mice (E3 open bars, E4 dotted bars). d) Spatial gene expression of *APOE, Gfap* and *Serpina3n* across the brain.

**Extended Data Fig. 7 (related to Fig. 7).**
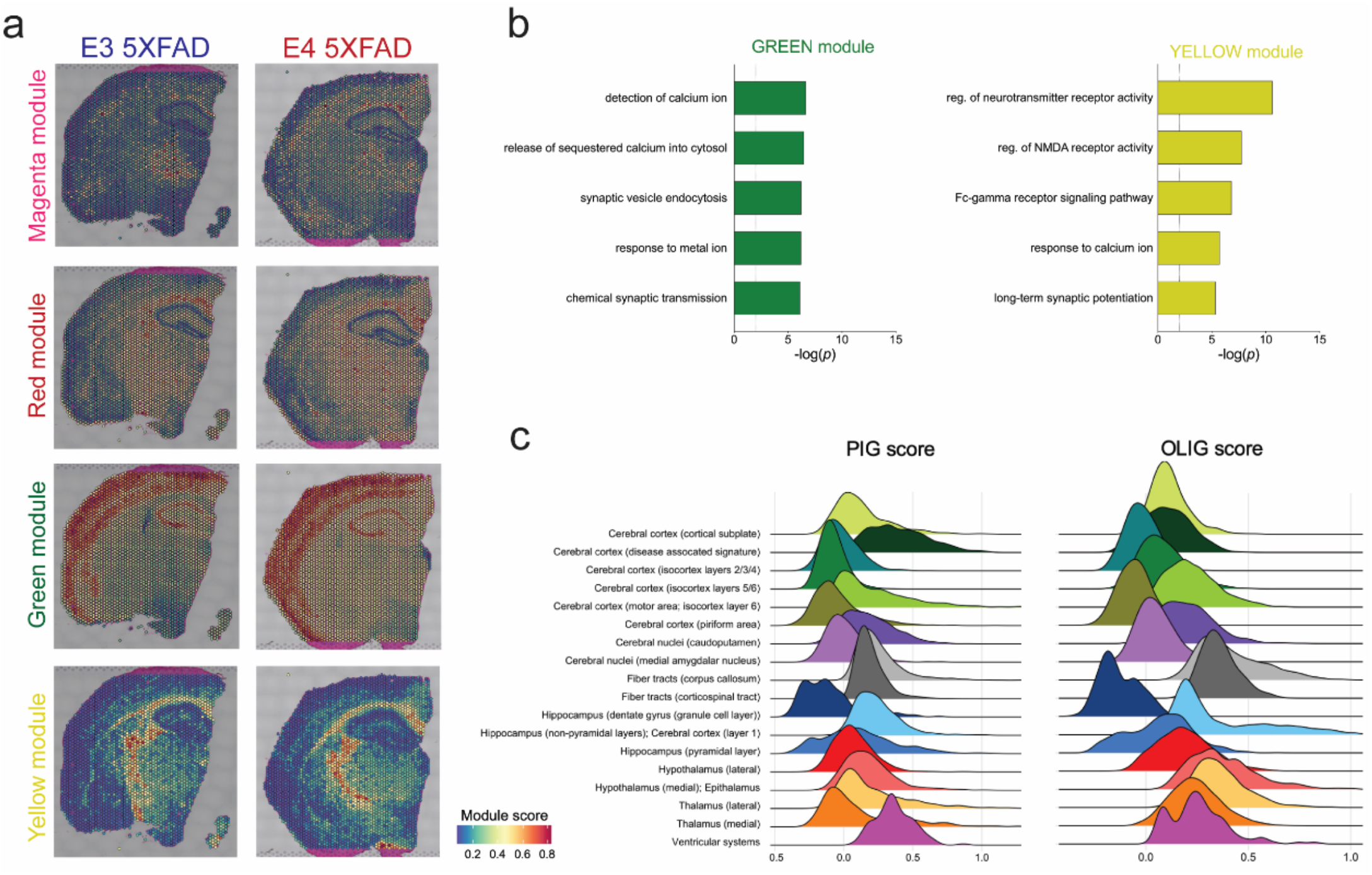
Plaque intensity module scores, gene ontology and PIG/OLIG module scores by brain region. a) Spatial expression of the four gene networks (modules) significantly associated with plaque intensity in E3 5XFAD (left) and E4 5XFAD (right) brains. b) Top 5 gene ontology terms associated with the two modules (green, yellow) negatively correlated with plaque intensity. c) Ridge plots showing PIG and OLIG module scores for each spatial transcriptomics cluster (with corresponding anatomical assignment). Module scores are composite scores for all 6 spatial transcriptomics brains combined.

**Extended Data Fig. 8 (related to Fig. 8).**
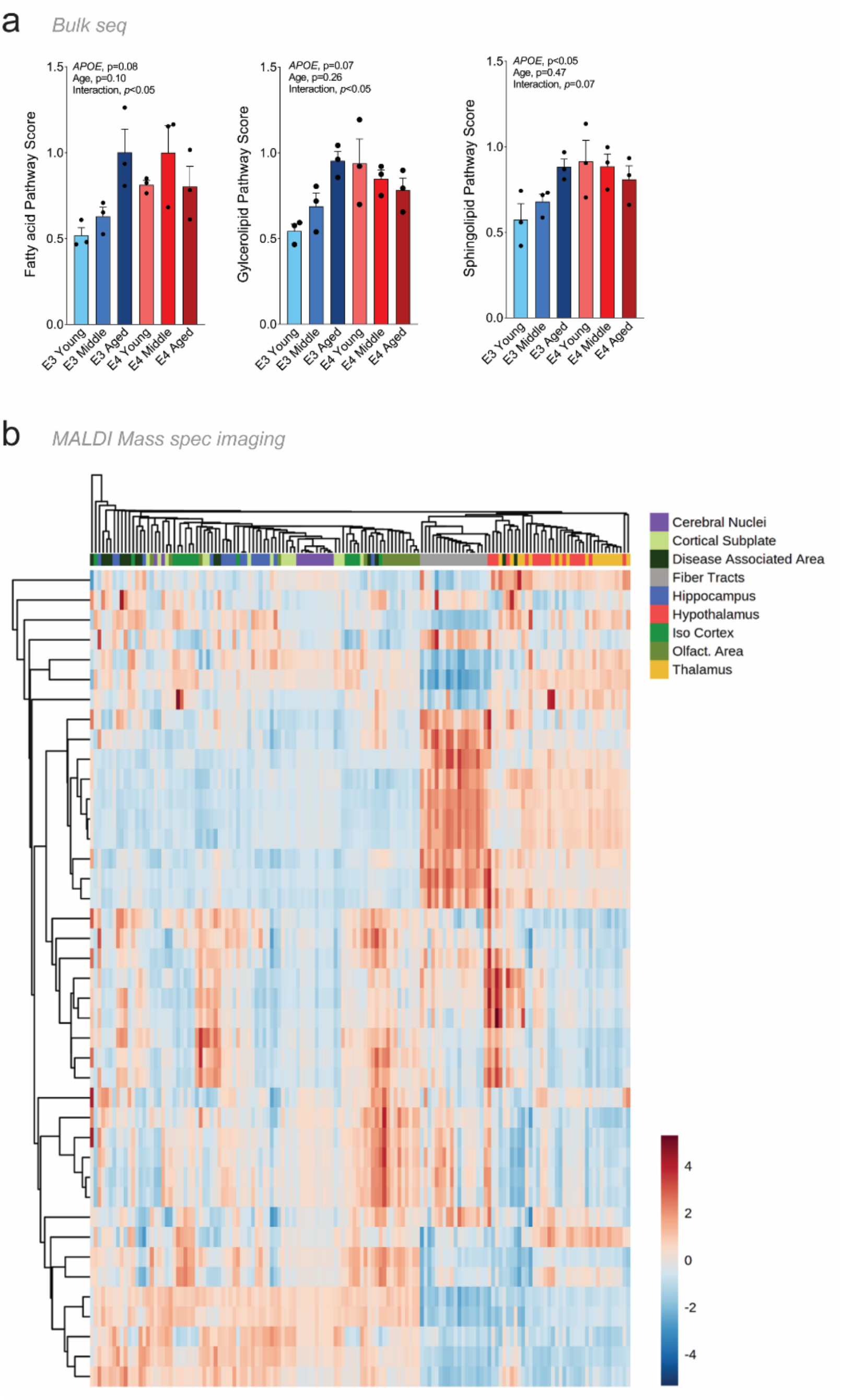
Lipid metabolism pathways from bulk sequencing, MALDI MSI quantification of lipids by brain region. **a**, Expression of fatty acid, glycerolipid, and sphingolipid pathway genes increase with *APOE* and/or age in whole brain tissue. **b**, Targeted lipidomics via MALDI MSI shows clear clustering by brain region, with the exception of the cortical and subcortical “disease associated area” found primarily in E4 5XFAD mice. Heat map displaying intensity of lipids (rows) quantified within each brain region (column). *n=3* mice per group.

## References

1 Jansen, I. E. et al. Genome-wide meta-analysis identifies new loci and functional pathways influencing Alzheimer’s disease risk. Nat Genet 51, 404–413, doi:10.1038/s41588-018-0311-9 (2019).

2 Kunkle, B. W. et al. Genetic meta-analysis of diagnosed Alzheimer’s disease identifies new risk loci and implicates Aβ, tau, immunity and lipid processing. Nat Genet 51, 414–430, doi:10.1038/s41588-019-0358-2 (2019).

3 Bertram, L., McQueen, M. B., Mullin, K., Blacker, D. & Tanzi, R. E. Systematic meta-analyses of Alzheimer disease genetic association studies: the AlzGene database. Nat Genet 39, 17–23, doi:10.1038/ng1934 (2007).

4 Bertram, L. & Tanzi, R. E. Genome-wide association studies in Alzheimer’s disease. Hum Mol Genet 18, R137–145, doi:10.1093/hmg/ddp406 (2009).

5 Harold, D. et al. Genome-wide association study identifies variants at CLU and PICALM associated with Alzheimer’s disease. Nat Genet 41, 1088–1093, doi:10.1038/ng.440 (2009).

6 Mosconi, L. Glucose metabolism in normal aging and Alzheimer’s disease: Methodological and physiological considerations for PET studies. Clinical and translational imaging 1, doi:10.1007/s40336-013-0026-y (2013).

7 Johnson, E. C. B. et al. Large-scale proteomic analysis of Alzheimer’s disease brain and cerebrospinal fluid reveals early changes in energy metabolism associated with microglia and astrocyte activation. Nature medicine, doi:10.1038/s41591-020-0815-6 (2020).

8 Oresic, M. et al. Metabolome in progression to Alzheimer’s disease. Transl Psychiatry 1, e57, doi:10.1038/tp.2011.55 (2011).

9 Xu, X. H., Huang, Y., Wang, G. & Chen, S. D. Metabolomics: a novel approach to identify potential diagnostic biomarkers and pathogenesis in Alzheimer’s disease. Neurosci Bull 28, 641–648, doi:10.1007/s12264-012-1272-0 (2012).

10 Arnold, M. et al. Sex and APOE ε4 genotype modify the Alzheimer’s disease serum metabolome. Nat Commun 11, 1148, doi:10.1038/s41467-020-14959-w (2020).

11 Devanney, N. A., Stewart, A. N. & Gensel, J. C. Microglia and macrophage metabolism in CNS injury and disease: The role of immunometabolism in neurodegeneration and neurotrauma. Exp Neurol 329, 113310, doi:10.1016/j.expneurol.2020.113310 (2020).

12 O’Neill, L. A., Kishton, R. J. & Rathmell, J. A guide to immunometabolism for immunologists. Nat Rev Immunol 16, 553–565, doi:10.1038/nri.2016.70 (2016).

13 Fernandez, C. G., Hamby, M. E., McReynolds, M. L. & Ray, W. J. The Role of APOE4 in Disrupting the Homeostatic Functions of Astrocytes and Microglia in Aging and Alzheimer’s Disease. Front Aging Neurosci 11, 14, doi:10.3389/fnagi.2019.00014 (2019).

14 Raber, J., Huang, Y. & Ashford, J. W. ApoE genotype accounts for the vast majority of AD risk and AD pathology. Neurobiol Aging 25, 641–650, doi:10.1016/j.neurobiolaging.2003.12.023 (2004).

15 Sala Frigerio, C. et al. The Major Risk Factors for Alzheimer’s Disease: Age, Sex, and Genes Modulate the Microglia Response to Abeta Plaques. Cell Rep 27, 1293–1306 e1296, doi:10.1016/j.celrep.2019.03.099 (2019).

16 Chen, W. T. et al. Spatial Transcriptomics and In Situ Sequencing to Study Alzheimer’s Disease. Cell, doi:10.1016/j.cell.2020.06.038 (2020).

17 Krasemann, S. et al. The TREM2-APOE Pathway Drives the Transcriptional Phenotype of Dysfunctional Microglia in Neurodegenerative Diseases. Immunity 47, 566–581 e569, doi:10.1016/j.immuni.2017.08.008 (2017).

18 Keren-Shaul, H. et al. A Unique Microglia Type Associated with Restricting Development of Alzheimer’s Disease. Cell 169, 1276–1290 e1217, doi:10.1016/j.cell.2017.05.018 (2017).

19 Olah, M. et al. A transcriptomic atlas of aged human microglia. Nat Commun 9, 539, doi:10.1038/s41467-018-02926-5 (2018).

20 Ping, L. et al. Global quantitative analysis of the human brain proteome in Alzheimer’s and Parkinson’s Disease. Sci Data 5, 180036, doi:10.1038/sdata.2018.36 (2018).

21 Rangaraju, S. et al. Quantitative proteomics of acutely-isolated mouse microglia identifies novel immune Alzheimer’s disease-related proteins. Molecular Neurodegeneration 13, 34, doi:10.1186/s13024-018-0266-4 (2018).

22 Hickman, S. E. et al. The microglial sensome revealed by direct RNA sequencing. Nat Neurosci 16, 1896–1905, doi:10.1038/nn.3554 (2013).

23 Orre, M. et al. Isolation of glia from Alzheimer’s mice reveals inflammation and dysfunction. Neurobiol Aging 35, 2746–2760, doi:10.1016/j.neurobiolaging.2014.06.004 (2014).

24 Wang, Y. et al. TREM2 lipid sensing sustains the microglial response in an Alzheimer’s disease model. Cell 160, 1061–1071, doi:10.1016/j.cell.2015.01.049 (2015).

25 Mathys, H. et al. Single-cell transcriptomic analysis of Alzheimer’s disease. Nature 570, 332–337, doi:10.1038/s41586-019-1195-2 (2019).

26 Del-Aguila, J. L. et al. A single-nuclei RNA sequencing study of Mendelian and sporadic AD in the human brain. Alzheimer’s research & therapy 11, 71, doi:10.1186/s13195-019-0524-x (2019).

27 Srinivasan, K. et al. Alzheimer’s Patient Microglia Exhibit Enhanced Aging and Unique Transcriptional Activation. Cell Rep 31, 107843, doi:10.1016/j.celrep.2020.107843 (2020).

28 Zhao, N. et al. Alzheimer’s Risk Factors Age, APOE Genotype, and Sex Drive Distinct Molecular Pathways. Neuron 106, 727–742.e726, doi:10.1016/j.neuron.2020.02.034 (2020).

29 Nuriel, T. et al. The Endosomal-Lysosomal Pathway Is Dysregulated by APOE4 Expression in Vivo. Front Neurosci 11, 702, doi:10.3389/fnins.2017.00702 (2017).

30 Area-Gomez, E. et al. APOE4 is Associated with Differential Regional Vulnerability to Bioenergetic Deficits in Aged APOE Mice. Sci Rep 10, 4277, doi:10.1038/s41598-020-61142-8 (2020).

31 Nuriel, T. et al. Neuronal hyperactivity due to loss of inhibitory tone in APOE4 mice lacking Alzheimer’s disease-like pathology. Nat Commun 8, 1464, doi:10.1038/s41467-017-01444-0 (2017).

32 Miranda, A. M. et al. Effects of APOE4 allelic dosage on lipidomic signatures in the entorhinal cortex of aged mice. Transl Psychiatry 12, 129, doi:10.1038/s41398-022-01881-6 (2022).

33 Serrano-Pozo, A. et al. Effect of APOE alleles on the glial transcriptome in normal aging and Alzheimer’s disease. Nature Aging 1, 919–931, doi:10.1038/s43587-021-00123-6 (2021).

34 Aibar, S. et al. SCENIC: single-cell regulatory network inference and clustering. Nat Methods 14, 1083-1086, doi:10.1038/nmeth.4463 (2017).

35 Song, P. et al. Frameshift mutation of Timm8a1 gene in mouse leads to an abnormal mitochondrial structure in the brain, correlating with hearing and memory impairment. J Med Genet 58, 619–627, doi:10.1136/jmedgenet-2020-106925 (2021).

36 Weber, L. W., Boll, M. & Stampfl, A. Maintaining cholesterol homeostasis: sterol regulatory elementbinding proteins. World J Gastroenterol 10, 3081–3087, doi:10.3748/wjg.v10.i21.3081 (2004).

37 March-Diaz, R. et al. Hypoxia compromises the mitochondrial metabolism of Alzheimer’s disease microglia via HIF1. Nature Aging 1, 385–399, doi:10.1038/s43587-021-00054-2 (2021).

38 Lampropoulou, V. et al. Itaconate Links Inhibition of Succinate Dehydrogenase with Macrophage Metabolic Remodeling and Regulation of Inflammation. Cell Metab 24, 158–166, doi:10.1016/j.cmet.2016.06.004 (2016).

39 Mills, E. L. et al. Itaconate is an anti-inflammatory metabolite that activates Nrf2 via alkylation of KEAP1. Nature 556, 113–117, doi:10.1038/nature25986 (2018).

40 Liao, S. T. et al. 4-Octyl itaconate inhibits aerobic glycolysis by targeting GAPDH to exert anti-inflammatory effects. Nat Commun 10, 5091, doi:10.1038/s41467-019-13078-5 (2019).

41 Viola, A., Munari, F., Sánchez-Rodríguez, R., Scolaro, T. & Castegna, A. The Metabolic Signature of Macrophage Responses. Front Immunol 10, 1462, doi:10.3389/fimmu.2019.01462 (2019).

42 Clarke, J. R., Ribeiro, F. C., Frozza, R. L., De Felice, F. G. & Lourenco, M. V. Metabolic Dysfunction in Alzheimer’s Disease: From Basic Neurobiology to Clinical Approaches. J Alzheimers Dis 64, S405–s426, doi:10.3233/jad-179911 (2018).

43 Kinney, J. W. et al. Inflammation as a central mechanism in Alzheimer’s disease. Alzheimers Dement (N Y) 4, 575–590, doi:10.1016/j.trci.2018.06.014 (2018).

44 Meraz-Ríos, M. A., Toral-Rios, D., Franco-Bocanegra, D., Villeda-Hernández, J. & Campos-Peña, V. Inflammatory process in Alzheimer’s Disease. Front Integr Neurosci 7, 59, doi:10.3389/fnint.2013.00059 (2013).

45 Xiang, X. et al. Microglial activation states drive glucose uptake and FDG-PET alterations in neurodegenerative diseases. Sci Transl Med 13, eabe5640, doi:10.1126/scitranslmed.abe5640 (2021).

46 Malik, M. et al. Genetics ignite focus on microglial inflammation in Alzheimer’s disease. Mol Neurodegener 10, 52, doi:10.1186/s13024-015-0048-1 (2015).

47 Karch, C. M. & Goate, A. M. Alzheimer’s disease risk genes and mechanisms of disease pathogenesis. Biol Psychiatry 77, 43–51, doi:10.1016/j.biopsych.2014.05.006 (2015).

48 Efthymiou, A. G. & Goate, A. M. Late onset Alzheimer’s disease genetics implicates microglial pathways in disease risk. Mol Neurodegener 12, 43, doi:10.1186/s13024-017-0184-x (2017).

49 Shippy, D. C. & Ulland, T. K. Microglial Immunometabolism in Alzheimer’s Disease. Front Cell Neurosci 14, 563446, doi:10.3389/fncel.2020.563446 (2020).

50 Ortiz-Barahona, A., Villar, D., Pescador, N., Amigo, J. & del Peso, L. Genome-wide identification of hypoxia-inducible factor binding sites and target genes by a probabilistic model integrating transcription-profiling data and in silico binding site prediction. Nucleic Acids Research 38, 2332–2345, doi:10.1093/nar/gkp1205 (2010).

51 Grubman, A. et al. Transcriptional signature in microglia associated with Aβ plaque phagocytosis. Nature Communications 12, 3015, doi:10.1038/s41467-021-23111-1 (2021).

52 Nguyen, A. T. et al. APOE and TREM2 regulate amyloid-responsive microglia in Alzheimer’s disease. Acta neuropathologica 140, 477–493, doi:10.1007/s00401-020-02200-3 (2020).

53 Gale, S. C. et al. APOε4 is associated with enhanced in vivo innate immune responses in human subjects. J Allergy Clin Immunol 134, 127–134, doi:10.1016/j.jaci.2014.01.032 (2014).

54 Vitek, M. P., Brown, C. M. & Colton, C. A. APOE genotype-specific differences in the innate immune response. Neurobiol Aging 30, 1350–1360, doi:10.1016/j.neurobiolaging.2007.11.014 (2009).

55 Konttinen, H. et al. PSEN1ΔE9, APPswe, and APOE4 Confer Disparate Phenotypes in Human iPSC-Derived Microglia. Stem Cell Reports 13, 669–683, doi:10.1016/j.stemcr.2019.08.004 (2019).

56 Navarro, J. F. et al. Spatial Transcriptomics Reveals Genes Associated with Dysregulated Mitochondrial Functions and Stress Signaling in Alzheimer Disease. iScience 23, 101556–101556, doi:10.1016/j.isci.2020.101556 (2020).

57 Brown, C. M. et al. Apolipoprotein E isoform mediated regulation of nitric oxide release. Free Radic Biol Med 32, 1071–1075, doi:10.1016/s0891-5849(02)00803-1 (2002).

58 Zhu, Y. et al. APOE genotype alters glial activation and loss of synaptic markers in mice. Glia 60, 559–569, doi:10.1002/glia.22289 (2012).

59 Yin, C. et al. ApoE attenuates unresolvable inflammation by complex formation with activated C1q. Nature medicine 25, 496–506, doi:10.1038/s41591-018-0336-8 (2019).

60 Chung, W. S. et al. Novel allele-dependent role for APOE in controlling the rate of synapse pruning by astrocytes. Proc Natl Acad Sci U S A 113, 10186–10191, doi:10.1073/pnas.1609896113 (2016).

61 Kosicek, M. & Hecimovic, S. Phospholipids and Alzheimer’s Disease: Alterations, Mechanisms and Potential Biomarkers. International journal of molecular sciences 14, 1310–1322 (2013).

62 Lefterov, I. et al. APOE2 orchestrated differences in transcriptomic and lipidomic profiles of postmortem AD brain. Alzheimer’s research & therapy 11, 113, doi:10.1186/s13195-019-0558-0 (2019).

63 Chang, R. et al. Predictive metabolic networks reveal sex-and APOE genotype-specific metabolic signatures and drivers for precision medicine in Alzheimer’s disease. Alzheimer’s & dementia : the journal of the Alzheimer’s Association, doi:10.1002/alz.12675 (2022).

64 Fitz, N. F. et al. Phospholipids of APOE lipoproteins activate microglia in an isoform-specific manner in preclinical models of Alzheimer’s disease. Nature Communications 12, 3416, doi:10.1038/s41467-021-23762-0 (2021).

65 Tian, Q., Mitchell, B. A., Corkum, A. E., Moaddel, R. & Ferrucci, L. Metabolites Associated with Memory and Gait: A Systematic Review. Metabolites 12, doi:10.3390/metabo12040356 (2022).

66 Chen, R., Feldstein, A. E. & McIntyre, T. M. Suppression of mitochondrial function by oxidatively truncated phospholipids is reversible, aided by bid, and suppressed by Bcl-XL. J Biol Chem 284, 26297–26308, doi:10.1074/jbc.M109.018978 (2009).

67 Bochkov, V. et al. Pleiotropic effects of oxidized phospholipids. Free Radic Biol Med 111, 6–24, doi:10.1016/j.freeradbiomed.2016.12.034 (2017).

68 Hazen, S. L. Oxidized phospholipids as endogenous pattern recognition ligands in innate immunity. J Biol Chem 283, 15527–15531, doi:10.1074/jbc.R700054200 (2008).

69 Binder, C. J., Papac-Milicevic, N. & Witztum, J. L. Innate sensing of oxidation-specific epitopes in health and disease. Nat Rev Immunol 16, 485–497, doi:10.1038/nri.2016.63 (2016).

70 Farmer, B. C., Walsh, A. E., Kluemper, J. C. & Johnson, L. A. Lipid Droplets in Neurodegenerative Disorders. Front Neurosci 14, 742, doi:10.3389/fnins.2020.00742 (2020).

71 Farmer, B. C., Kluemper, J. & Johnson, L. A. Apolipoprotein E4 Alters Astrocyte Fatty Acid Metabolism and Lipid Droplet Formation. Cells 8, doi:10.3390/cells8020182 (2019).

72 Marschallinger, J. et al. Lipid-droplet-accumulating microglia represent a dysfunctional and proinflammatory state in the aging brain. Nat Neurosci 23, 194–208, doi:10.1038/s41593-019-0566-1 (2020).

73 Sienski, G. et al. APOE4 disrupts intracellular lipid homeostasis in human iPSC-derived glia. Sci Transl Med 13, doi:10.1126/scitranslmed.aaz4564 (2021).

74 Machlovi, S. I. et al. APOE4 confers transcriptomic and functional alterations to primary mouse microglia. Neurobiol Dis 164, 105615, doi:10.1016/j.nbd.2022.105615 (2022).

75 Lin, Y. T. et al. APOE4 Causes Widespread Molecular and Cellular Alterations Associated with Alzheimer’s Disease Phenotypes in Human iPSC-Derived Brain Cell Types. Neuron 98, 1141–1154.e1147, doi:10.1016/j.neuron.2018.05.008 (2018).

76 Qi, G. et al. ApoE4 Impairs Neuron-Astrocyte Coupling of Fatty Acid Metabolism. Cell Rep 34, 108572, doi:10.1016/j.celrep.2020.108572 (2021).

77 Mahley, R. W., Weisgraber, K. H. & Huang, Y. Apolipoprotein E: structure determines function, from atherosclerosis to Alzheimer’s disease to AIDS. Journal of lipid research 50, S183–S188, doi:https://doi.org/10.1194/jlr.R800069-JLR200 (2009).

78 Saito, H. et al. Effects of Polymorphism on the Lipid Interaction of Human Apolipoprotein E*. Journal of Biological Chemistry 278, 40723–40729, doi:https://doi.org/10.1074/jbc.M304814200 (2003).

79 Sullivan, P. M. et al. Targeted replacement of the mouse apolipoprotein E gene with the common human APOE3 allele enhances diet-induced hypercholesterolemia and atherosclerosis. J Biol Chem 272, 17972–17980 (1997).

80 Knouff, C. et al. Apo E structure determines VLDL clearance and atherosclerosis risk in mice. J Clin Invest 103, 1579–1586, doi:10.1172/JCI6172 (1999).

81 Sullivan, P. M., Mezdour, H., Quarfordt, S. H. & Maeda, N. Type III hyperlipoproteinemia and spontaneous atherosclerosis in mice resulting from gene replacement of mouse Apoe with human Apoe*2. J Clin Invest 102, 130–135, doi:10.1172/JCI2673 (1998).

82 Sullivan, P. M., Mace, B. E., Maeda, N. & Schmechel, D. E. Marked regional differences of brain human apolipoprotein E expression in targeted replacement mice. Neuroscience 124, 725–733, doi:10.1016/j.neuroscience.2003.10.011 (2004).

83 Patel, T. et al. Transcriptional landscape of human microglia implicates age, sex, and APOE-related immunometabolic pathway perturbations. Aging Cell, e13606, doi:10.1111/acel.13606 (2022).

84 Johnson, E. C. B. et al. Large-scale deep multi-layer analysis of Alzheimer’s disease brain reveals strong proteomic disease-related changes not observed at the RNA level. Nat Neurosci 25, 213–225, doi:10.1038/s41593-021-00999-y (2022).

85 Dai, J. et al. Effects of APOE Genotype on Brain Proteomic Network and Cell Type Changes in Alzheimer’s Disease. Front Mol Neurosci 11, 454, doi:10.3389/fnmol.2018.00454 (2018).

86 Konijnenberg, E. et al. APOE ε4 genotype-dependent cerebrospinal fluid proteomic signatures in Alzheimer’s disease. Alzheimer’s research & therapy 12, 65, doi:10.1186/s13195-020-00628-z (2020).

87 Shuken, S. R. et al. Limited proteolysis–mass spectrometry reveals aging-associated changes in cerebrospinal fluid protein abundances and structures. Nature Aging, doi:10.1038/s43587-022-00196-x (2022).

88 Kaur, G., Poljak, A., Masters, C. L., Fowler, C. & Sachdev, P. Impact of APOE ε3 and ε4 genotypes on plasma proteome signatures in Alzheimer’s disease. bioRxiv, 2022.2001.2029.478291, doi:10.1101/2022.01.29.478291 (2022).

89 Flowers, A., Bell-Temin, H., Jalloh, A., Stevens, S. M. & Bickford, P. C. Proteomic analysis of aged microglia: shifts in transcription, bioenergetics, and nutrient response. Journal of Neuroinflammation 14, 96, doi:10.1186/s12974-017-0840-7 (2017).

90 Ulland, T. K. et al. TREM2 Maintains Microglial Metabolic Fitness in Alzheimer’s Disease. Cell 170, 649–663.e613, doi:10.1016/j.cell.2017.07.023 (2017).

91 De Miguel, Z. et al. Exercise plasma boosts memory and dampens brain inflammation via clusterin. Nature 600, 494–499, doi:10.1038/s41586-021-04183-x (2021).

92 Ha, J. et al. Plasma Clusterin as a Potential Link Between Diabetes and Alzheimer Disease. The Journal of Clinical Endocrinology & Metabolism 105, 3058–3068, doi:10.1210/clinem/dgaa378 (2020).

93 Zafar, A., Ng, H. P., Kim, G. D., Chan, E. R. & Mahabeleshwar, G. H. BHLHE40 promotes macrophage pro-inflammatory gene expression and functions. Faseb j 35, e21940, doi:10.1096/fj.202100944R (2021).

94 Pang, Z. et al. MetaboAnalyst 5.0: narrowing the gap between raw spectra and functional insights. Nucleic Acids Research 49, W388–W396, doi:10.1093/nar/gkab382 (2021).

95 Early, A. N., Gorman, A. A., Van Eldik, L. J., Bachstetter, A. D. & Morganti, J. M. Effects of advanced age upon astrocyte-specific responses to acute traumatic brain injury in mice. J Neuroinflammation 17, 115, doi:10.1186/s12974-020-01800-w (2020).

96 McKenzie, A. T. et al. Brain Cell Type Specific Gene Expression and Co-expression Network Architectures. Scientific Reports 8, 8868, doi:10.1038/s41598-018-27293-5 (2018).

97 Zeisel, A. et al. Molecular Architecture of the Mouse Nervous System. Cell 174, 999–1014.e1022, doi:https://doi.org/10.1016/j.cell.2018.06.021 (2018).

98 Ximerakis, M. et al. Single-cell transcriptomic profiling of the aging mouse brain. Nat Neurosci 22, 1696–1708, doi:10.1038/s41593-019-0491-3 (2019).

99 Han, X. et al. Mapping the Mouse Cell Atlas by Microwell-Seq. Cell 172, 1091–1107.e1017, doi:https://doi.org/10.1016/j.cell.2018.02.001 (2018).

100 Kuleshov, M. V. et al. Enrichr: a comprehensive gene set enrichment analysis web server 2016 update. Nucleic Acids Research 44, W90–W97, doi:10.1093/nar/gkw377 (2016).

101 Slenter, D. N. et al. WikiPathways: a multifaceted pathway database bridging metabolomics to other omics research. Nucleic Acids Res 46, D661–d667, doi:10.1093/nar/gkx1064 (2018).

102 Langfelder, P. & Horvath, S. WGCNA: an R package for weighted correlation network analysis. BMC Bioinformatics 9, 559, doi:10.1186/1471-2105-9-559 (2008).

103 Van de Sande, B. et al. A scalable SCENIC workflow for single-cell gene regulatory network analysis. Nature Protocols 15, 2247–2276, doi:10.1038/s41596-020-0336-2 (2020).

104 Lambert, S. A. et al. The Human Transcription Factors. Cell 172, 650–665, doi:10.1016/j.cell.2018.01.029 (2018).

105 Palla, G. et al. Squidpy: a scalable framework for spatial single cell analysis. bioRxiv (2021).

106 Jin, S. et al. Inference and analysis of cell-cell communication using CellChat. Nature Communications 12, 1088, doi:10.1038/s41467-021-21246-9 (2021).

107 Sheikh, B. N. et al. Systematic Identification of Cell-Cell Communication Networks in the Developing Brain. iScience 21, 273–287, doi:10.1016/j.isci.2019.10.026 (2019).

108 Hawkinson, T. R. et al. In situ spatial glycomic imaging of mouse and human Alzheimer’s disease brains. Alzheimer’s & dementia : the journal of the Alzheimer’s Association, doi:10.1002/alz.12523 (2021).

